# Heterosynaptic Plasticity of the Visuo-auditory Projection Requires Cholecystokinin released from Entorhinal Cortex Afferents

**DOI:** 10.1101/2022.10.04.510820

**Authors:** Wenjian Sun, Haohao Wu, Yujie Peng, Peng Tang, Ming Zhao, Xuejiao Zheng, Hemin Feng, Hao Li, Ye Liang, Jing Li, Junfeng Su, Xi Chen, Tomas Hökfelt, Jufang He

**Affiliations:** Department of Neuroscience, City University of Hong Kong, Kowloon Tong, Hong Kong SAR, P.R. China, 0000; Centre for Regenerative Medicine and Health, Hong Kong Institute of Science & Innovation, Chinese Academy of Sciences, Hong Kong SAR, P.R. China, 0000; Department of Neuroscience, Karolinska Institutet, Stockholm, Sweden, 17177; City University of Hong Kong Shenzhen Research Institute, Shenzhen, China, 518507; Institute of Advanced Study, City University of Hong Kong, Kowloon Tong, Hong Kong SAR, P.R. China, 0000; Zilkha Neurogenetic Institute, University of Southern California, Los Angeles, CA, USA, 90033; Biozentrum, Department of Cell Biology, University of Basel, Basel, Switzerland, CH-4046; Friedrich Miescher Institute for Biomedical Research, Basel, Switzerland, CH-4058; Beijing Genomics Institute-Shenzhen, Shenzhen, China, 518103; Department of Neurosurgery, Stanford University School of Medicine, Stanford, CA, USA, 94305; F.M. Kirby Neurobiology Center, Boston Children’s Hospital, Boston, MA, USA, 02115

**Keywords:** auditory cortex, cross-modal association, Hebbian plasticity, long-term potentiation, neuropeptides, optogenetics, pre- and post-synaptic coactivity, transmitter coexistence, visuo-auditory memory

## Abstract

The entorhinal cortex is involved in establishing enduring visuo-auditory associative memory in the neocortex. Here we explored the mechanisms underlying this synaptic plasticity related to projections from the visual and entorhinal cortices to the auditory cortex, using optogenetics of dual pathways. High-frequency laser stimulation (HFLS) of the visuo-auditory projection did not induce long-term potentiation (LTP). However, after pairing with sound stimulus, the visuo-auditory inputs were potentiated following either infusion of cholecystokinin (CCK) or HFLS of the entorhino-auditory CCK-expressing projection. Combining retrograde tracing and RNAscope in situ hybridization, we show that CCK expression is higher in entorhinal cortex neurons projecting to the auditory cortex than in those originating from the visual cortex. In the presence of CCK, potentiation in the neocortex occurred when the pre-synaptic input arrived 200 ms before post-synaptic firing, even after just five trials of pairing. Behaviorally, inhibition of CCK signaling blocked the generation of associative memory. Our results indicate that neocortical visuo-auditory association is formed through heterosynaptic plasticity, which depends on release of CCK in the neocortex mostly from entorhinal afferents.

## Introduction

Cross-modal association is crucial for our brain to integrate information from different modalities to provide a useful output. Traditionally, this process is assumed to mainly occur in higher association cortices as evidenced by both anatomical(Cusick et al., 1995; Seltzer et al., 1996) and physiological(Fuster et al., 2000; Lipton et al., 1999; Sakai and Miyashita, 1991; Schlack et al., 2005; Sugihara et al., 2006) studies. Moreover, fMRI studies(Calvert et al., 1997; Finney et al., 2001; Foxe et al., 2002; Pekkola et al., 2006) and *in vivo* electrophysiological recordings(Brosch et al., 2005; Zhou and Fuster, 2004; Zhou and Fuster, 2000) have provided further evidence for involvement of unimodal sensory cortices. We have shown that neurons in the auditory cortex (AC) start to respond to light stimuli after classical fear conditioning, thus coupling light and electrical stimulation of the AC (ESAC) (Chen et al., 2013). These results indicate that the AC participates in the establishment of visuo-auditory association. However, the direct source of the visual signal to the AC is unclear. Here, we provide an anatomical foundation for a direct projection from the visual cortex (VC) to the AC by combining retrograde tracing and optogenetics in mice.

Observations of patient H.M. demonstrate that removal of the bilateral medial temporal lobe prevents the formation of long-term declarative memory(Milner and Klein, 2016; Scoville and Milner, 1957), suggesting that this lobe plays a crucial role in forming new memories about facts and events. The entorhinal cortex (EC) of the medial temporal lobe is strongly and reciprocally connected with both the neocortex and hippocampus(Canto et al., 2008; Swanson and Köhler, 1986).

It is well established that neocortex expresses a variety of neuropeptides, primarily in GABAergic, inhibitory interneurons (Hendry et al., 1984; Somogyi et al., 1984; Somogyi and Klausberger, 2005). The most abundant neuropeptide in brain is sulphated cholecystokinin octapeptide (CCK-8S) (Beinfeld et al., 1981; Dockray et al., 1978; Innis et al., 1979; Larsson and Rehfeld, 1979; Rehfeld, 1978; Vanderhaeghen et al., 1980), and this peptide was early shown to have excitatory effects on pyramidal neurons(Dodd and Kelly, 1981). CCK is, however, also found in pyramidal projection neurons, which have high levels of CCK transcript as shown with in situ hybridization (Burgunder and Young, 1988; Schiffmann and Vanderhaeghen, 1991; Siegel and Young, 1985). Several studies have reported that the EC is rich in CCK-positive ^(+)^ neurons, both in rat(Greenwood et al., 1981; Innis et al., 1979; Kohler and Chan-Palay, 1982) and mouse (Meziane et al., 1997). In fact, CCK plays an important role in learning and memory (Horinouchi et al., 2004; Lo et al., 2008; Meziane et al., 1993; Nomoto et al., 1999; Tsutsumi et al., 1999).

We have previously shown that the cortical projection neurons in the EC of mouse and rat mostly are CCK^+^ and glutamatergic, and that this pathway is important for visuo-auditory association (Chen et al., 2019; Li et al., 2014; Zhang et al., 2020). Briefly, this type of association can be blocked by inactivation of the EC (Chen et al., 2013) or local infusion of a CCK-B receptor (CCKBR) antagonist(Li et al., 2014). Local infusion of a CCK agonist enabled visual responses in the auditory cortex after paring a light stimulus with a noise burst stimulus(Li et al., 2014). Moreover, we presented evidence that CCK released from the EC projection in the AC enables LTP and sound-sound association (Chen et al., 2019). Thus, the CCKBR antagonist blocked HFLS-induced neocortical LTP, and CCK^-/-^ mice lacked such LTP (Chen et al., 2019). However, also an NMDAR antagonist blocked the LTP, a possible explanation being that CCK release is controlled by NMDA receptors (Chen et al., 2019).

In the present study we expanded our analysis of visuo-auditory association using two channelrhodopsins, Chronos and ChrimsonR, examining various ways to potentiate the responses to inputs/projection from VC to AC (VC→AC): (i) HFLS of VC→AC projection with classical high frequency stimulation protocol; (ii) infusion of a CCK agonist in the AC followed by pairing of presynaptic activation of VC→AC terminals expressing opsin by single pulse laser stimulation (SPLS) with postsynaptic noise-induced AC firing (Pre/Post Pairing); (iii) pairing of HFLS of EC→AC CCK^+^ projection with SPLS of the VC→AC projection; (iv) HFLS of EC→AC CCK^+^ projection followed by Pre/Post Pairing; (v) HFLS of VC→AC projection followed by Pre/Post Pairing; (vi) testing different parameters of the pairing protocol: the frequency of the laser stimulation of the EC→AC CCK^+^ projection; the delay between the termination of HFLS of EC→AC CCK^+^ projection and Pre/Post Pairing (Delay 1); and the delay between presynaptic activation of VC→AC projection and postsynaptic auditory cortex activation (Delay 2). Of particular interest was testing spike timing-dependent plasticity (STDP), an extension of the Hebbian learning rule, stating: in order to induce potentiation, the critical window between arrival of presynaptic input and postsynaptic firing should not be more than 20 ms (Bi and Poo, 1998; Markram et al., 1997; Zhang et al., 1998). This theory has been challenged(Bittner et al., 2017; Drew and Abbott, 2006; Izhikevich, 2007), and we hypothesized that endogenous CCK could be involved in a type of synaptic potentiation that is different from STDP. Secondly, in behavioral experiments we examined, if infusion of a CCK *antagonist* in the AC could block the visuo-auditory association, and if a CCK *agonist* could rescue the deficit of visuo-auditory association in CCK^-/-^ mice. Our results further support an important role of CCK for associative memory under natural conditions. Finally, we combined retrograde tracing and RNAscope in situ hybridization to explore CCK expression levels in the EC neurons projecting to the AC versus the projection from the VC.

## Results

### The auditory cortex receives a direct projection from the visual cortex

To examine the origin of the visual information underlying the previously observed visual responses in the AC(Chen et al., 2013; Li et al., 2014), we injected the retrograde tracer cholera toxin subunit B (Alexa Fluor 488 Conjugate, Molecular Probes, US) in the AC. The auditory thalamus was strongly labeled in the dorsal (MGD), ventral (MGV), and medial (MGM) subdivisions (Figure 1A1, 2). Many retrogradely labeled neurons were observed in both the primary and associative VC (Figure 1A3, 4), and were mostly distributed in layer V (Figure 1A4). The result here is consistent with previous studies reporting the existence of reciprocal projections between the VC and AC (Bizley et al., 2007; Budinger et al., 2006; Falchier et al., 2002; Falchier et al., 2010; Rockland and Ojima, 2003), and provides a possible anatomical basis for visuo-auditory associations formed in the AC.

**Figure 1.**
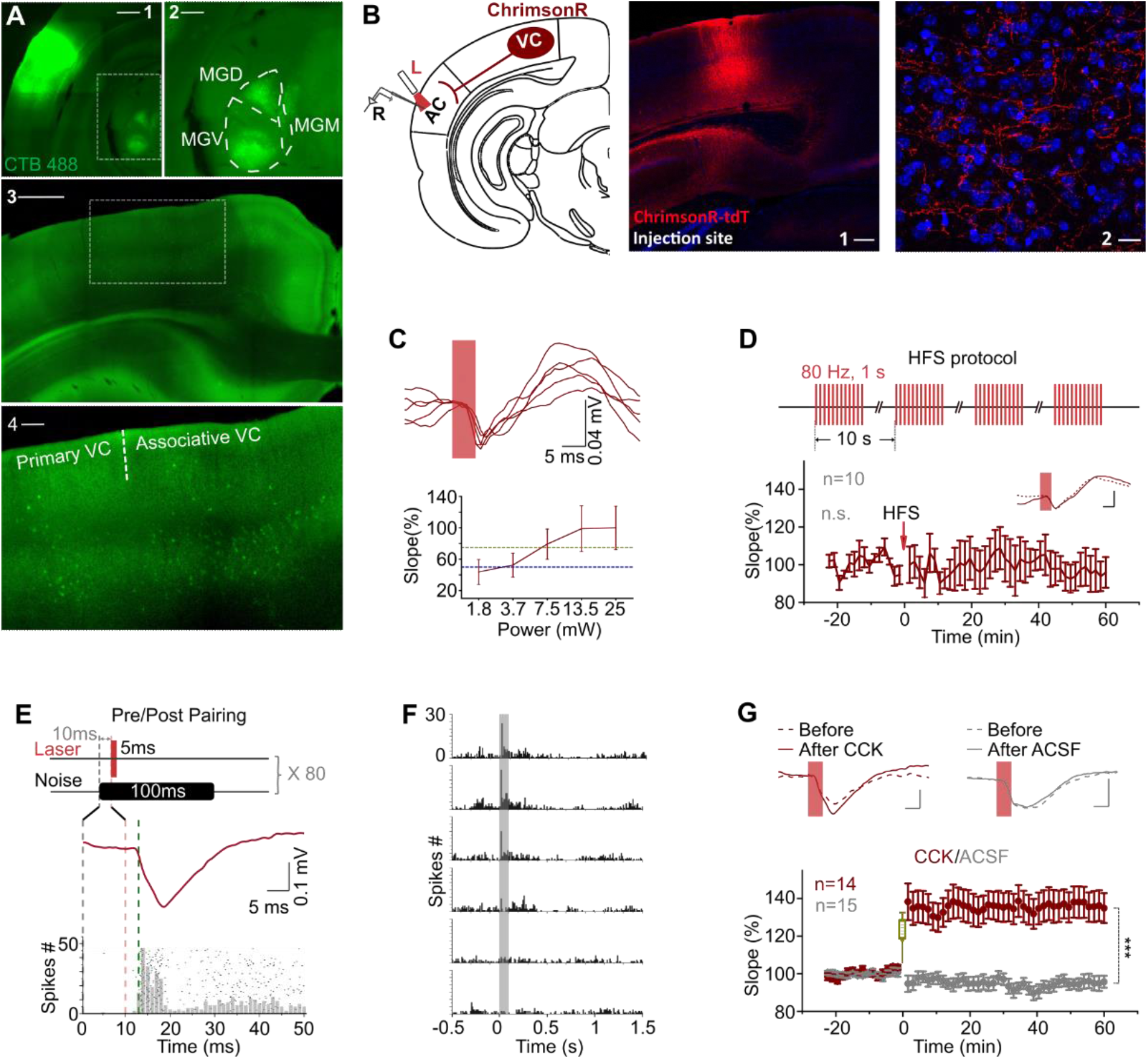
VC→AC projection was potentiated after pairing VALS evoked presynaptic activation with noise induced postsynaptic firing in the presence of CCK. (A) Images show the injection site of the retrograde tracer (CTB488) in the AC (A_1_, scale bar: 1000 μm) and retrogradely labeled neurons in the auditory thalamus (A_2_, an enlargement of the boxed area in the A_1_, scale bar: 500 μm) and the VC (A_3_, scale bar: 1000 μm; A_4_, an enlargement of the boxed area in A_3_, scale bar: 200 μm). MGV, MGD, and MGM are abbreviations for the ventral, dorsal, and medial parts of the medial geniculate nucleus, respectively. (B) Left: schematic drawing of the experimental setup. AAV9-syn-ChrimsonR-tdtomato was injected in the VC. L, laser fiber; R, recording electrode. Right: representative images of the injection site in the VC (1) and the projection terminals in the AC (2). Scale bars: 1, 200 μm; 2, 20 μm. (C) Example fEPSP traces evoked by laser stimulation at different intensities (upper) and the corresponding input/output curve (bottom). Blue and yellow dashed line indicate 50% and 75% of fEPSP saturation. (D) Upper: modified HFS protocol. Bottom: normalized fEPSP_VALS_ slopes before and after HFLS of VC→AC projection alone; inset, example traces before (dashed) and after (solid) HFLS. Error bars represent SEM. paired t-test, t (9) = 0.878, n.s. p = 0.403, n = 10. (E) The protocol of pairing VALS evoked presynaptic input with noise induced postsynaptic firing. (F) PSTHs of spike responses to noises at different sound intensities (40 to 90 dB, from bottom to top). (G) Normalized fEPSP_VALS_ slopes (bottom) and example traces (upper) before and after Pre/Post Pairing with CCK-8S (red) or ACSF (gray) infusion in the AC. Error bars represent SEM. *** p < 0.001, n = 14 for CCK group, n = 15 for ACSF group, two-way RM ANOVA with Bonferroni post-hoc test. See Table S1 for detailed Statistics.

### High-frequency stimulation of VC→AC projection does not induce LTP

High-frequency stimulation (HFS) is a classical protocol to induce LTP (Bashir et al., 1991; Bliss and Gardner-Medwin, 1973; Bliss and Lomo, 1973; Hernandez et al., 2005; Yun et al., 2002), which typically consists of 1 s train of pulses at 100 Hz repeated 3 times with an inter-trail interval (ITI) of 10 s. Based on the current understanding we should be able to induce LTP, if the VC→AC projection is activated with HFLS. We injected AAV9-syn-ChrimsonR-tdtomato in the VC of wildtype mice and manipulated the VC projection terminals in the AC expressing ChimsonR (Figures 1B), a variant of channelrhodopsin-2. Laser stimulation of the VC→AC projection (VALS) induced a field excitatory post-synaptic potential (fEPSPVALS) in the AC as an indicator of the VC→AC input. To prevent photoelectric artifacts, fEPSPs evoked by laser stimulation were recorded by glass pipette electrodes with an impedance of 1 MΩ rather than metal electrodes(Cardin et al., 2010; Kozai and Vazquez, 2015) (Figures S1A and S1B). Generally, a laser with higher intensity induced a fEPSP with a steeper slope and larger amplitude until saturation was reached (Figure 1C). Considering the kinetics of ChrimsonR (Klapoetke et al., 2014), we modified the HFS protocol and used 4 trials of 1 s pulse train at 80 Hz with an ITI of 10 s (Figure 1D upper). We chose the laser intensity that induced a 50% fEPSP saturation for baseline and the post-HFLS tests and 75% for HFLS. However, no significant LTP was induced in the VC→AC projection by HFLS of this pathway alone (Figure 1D bottom, p = 0.403, n = 10, paired t-test).

### VC→AC inputs are potentiated after pairing the activation of their terminals with noise bursts in the presence of CCK

Hebbian theory says that cells that fire together wire together. We next tested if VC→AC inputs can be potentiated after pairing with repetitive AC activation. We used VALS to evoke presynaptic input and noise stimulus to trigger postsynaptic AC firing. Since the latency of fEPSP_VALS_ is approximately 2–2.5 ms, and the firing latency of noise responses in the AC of mice is mostly equal to or longer than 13 ms, we presented the laser stimulus 10 ms after noise. Therefore, we started the presynaptic input just before the postsynaptic firing (Pre/Post Pairing, Figure 1E). Responses to noise at different sound intensities were first tested (Figure 1F), and we chose the intensity that evoked reliable firing for Pre/Post Pairing. However, even after 80 trials of Pre/Post Pairing, the VC→AC inputs were not potentiated (Figure 1G, gray, ACSF group).

In the previous studies we have shown that CCK has an important role in neocortical plasticity (Chen et al., 2019; Li et al., 2014; Zhang et al., 2020). We then examined if VC→AC inputs could be potentiated after Pre/Post Pairing in the presence of CCK. Before Pre/Post Pairing, CCK-8S (10 ng/μL, 0.5 μL, 0.1 μL/min; Tocris Bioscience, Bristol, UK) was infused in the AC. In line with our hypothesis, VC→AC inputs were strongly potentiated after infusion of CCK-8S compared with ACSF. The averaged fEPSP_VALS_ slope increased immediately after pairing and remained elevated for 1 h in the CCK injection group (Figures 1G and S1C, two-way repeated measures [RM] ANOVA, F _(1,27)_ = 25.125, significant interaction, p < 0.001; red, CCK before [101.3±0.8%] vs. CCK after [136.5±7.7%], pairwise comparison, p < 0.001, n = 14; gray, ACSF before [99.9±0.8%] vs. ACSF after [95.3±2.9%], pairwise comparison, p = 0.411, n = 15). Collectively, these results provide evidence that CCK together with noise, but not noise alone, enables a visuo-auditory association in the AC via a direct projection from the VC to AC.

### HFLS of EC→AC CCK+ terminals results in LTP of VC→AC inputs after pairing with postsynaptic firing in the AC evoked by noise stimuli

We have shown that cortical projection neurons in the EC mostly are CCK^+^ and glutamatergic, and that HFS induces CCK release in the auditory cortex (Chen et al., 2019). We then explored if endogenous CCK could enable the potentiation of VC→AC inputs. With Chronos and ChrimsonR (Klapoetke et al., 2014), we were able to manipulate two different neural pathways with two color activation. We injected AAV9-Ef1α-Flex-Chronos-GFP and AAV9-hSyn-ChrimsonR-tdTomato in the EC and VC of the CCK-iRES-Cre mouse to activate the EC→AC CCK^+^ projection and VC→AC projection, respectively (Figure 2A). HFLS (473 nm 80 Hz, 5- ms/pulse, 10 pulses, Figure S2C left) of the EC→AC CCK^+^ projection was applied and, after a 10-ms interval, pairing of VALS (635 nm) and noise stimulus was followed (Figure 2C, HFLSEA/Pre/PostPairing). This protocol was repeated for 5 trials with an ITI of 10s. Likewise, we injected AAV9-Ef1α-Flex-Chronos-GFP in the VC of CaMKIIa-Cre mice (Figure 2B) and applied 5 trials of HFLS of VC→AC projection (Figure S2C right) followed by the pairing of VALS and noise stimulus (Figure 2D, HFLSVA/Pre/PostPairing).

**Figure 2.**
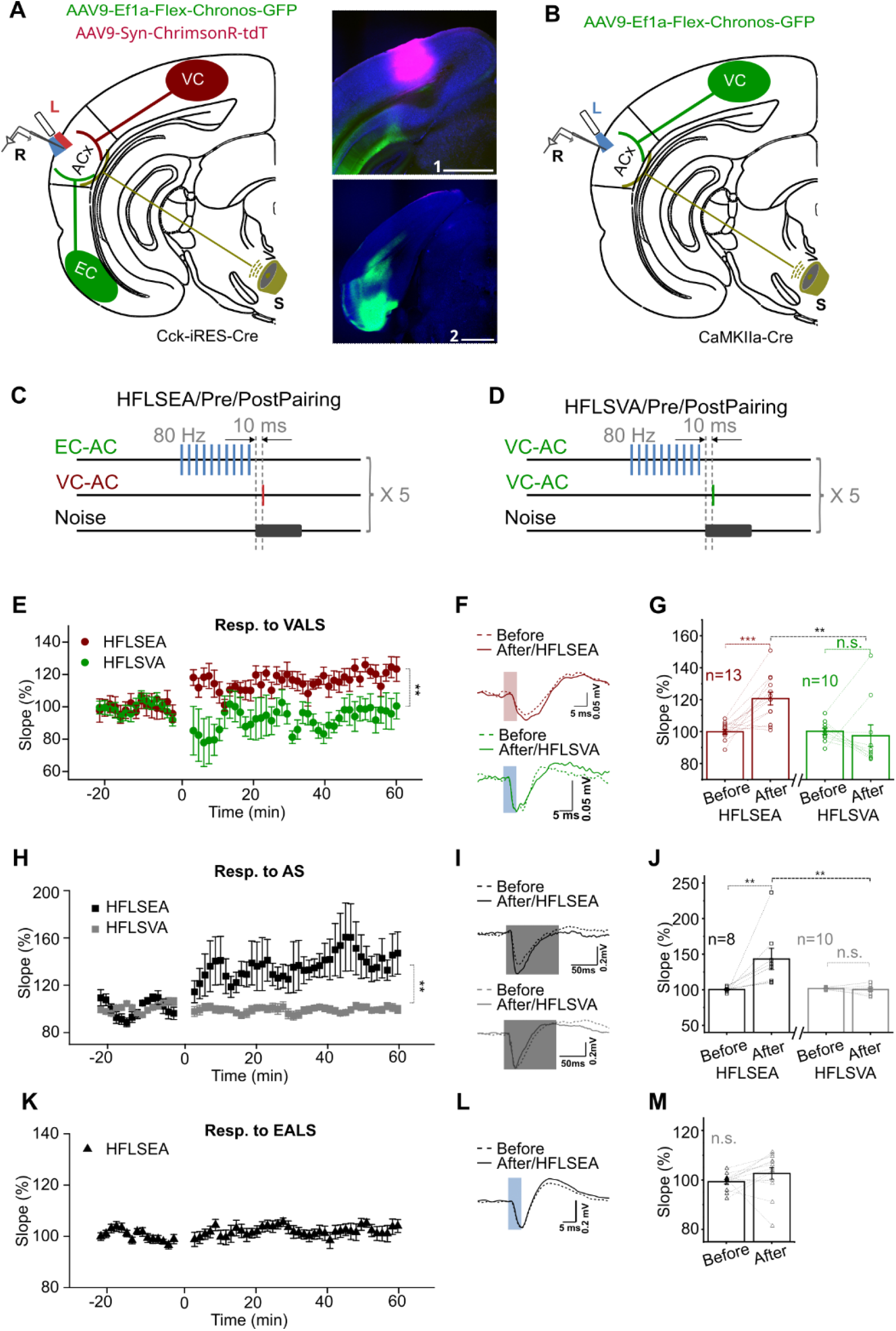
HFLS of EC→AC CCK^+^ projection but not VC→AC projection induced the potentiation of VC→AC inputs after pairing with noise evoked postsynaptic activation. (A) Left: schematic drawing of the experimental setup. AAV9-Ef1α-Flex- Chronos-GFP and AAV9-Syn-ChrimsonR-tdTomato were injected in the EC and VC of CCK-iRES-Cre mice respectively. Right: representative images of the injection sites in the VC (1) and the EC (2). Scale bars: 1, 1000 μm; 2, 1000 μm. L, laser fiber; R, recording electrode; S, sound. (B) Schematic drawing of the experimental setup. AAV9-Ef1α-Flex-Chronos-GFP was injected in the VC of CaMKIIa-cre mice. L, laser fiber; R, recording electrode; S, sound. (C) and (D) Protocols of HFLSEA/Pre/PostPairing and HFLSVA/Pre/PostPairing, respectively. (E) Normalized fEPSP_VALS_ slopes before and after HFLSEA/Pre/PostPairing (red) or HFLSVA/Pre/PostPairing (green). ** p < 0.01, two-way RM ANOVA with post-hoc Bonferroni test. (F) Example fEPSP_VALS_ traces before and after HFLSEA/Pre/PostPairing (red) or HFLSVA/Pre/PostPairing (green). Scale bars: upper, 5 ms and 0.05 mV; bottom, 5 ms and 0.05 mV. (G) Individual and average fEPSP_VALS_ slope changes before and after HFLSEA/Pre/PostPairing (red) or HFLSVA/Pre/PostPairing (green). ** p < 0.01, *** p < 0.001, n.s. p = 0.623, n = 13 for HFLSEA/Pre/PostPairing group, n = 10 for HFLSVA/Pre/PostPairing group, two-way RM ANOVA with post-hoc Bonferroni test. (H) Normalized fEPSP_AS_ slopes before and after HFLSEA/Pre/PostPairing (black) or HFLSVA/Pre/PostPairing (gray). ** p < 0.01, two-way RM ANOVA with post-hoc Bonferroni test. (I) Example fEPSP_AS_ traces before and after HFLSEA/Pre/PostPairing (black) or HFLSVA/Pre/PostPairing (gray). Scale bars: upper, 50 ms and 0.2 mV; bottom, 50 ms and 0.2 mV. (J) Individual and average fEPSP_AS_ slope changes before and after HFLSEA/Pre/PostPairing (black) or HFLSVA/Pre/PostPairing (gray). **p < 0.01, n.s. p = 0.898, n = 8 for HFLSEA/Pre/PostPairing group, n = 10 for HFLSVA/Pre/PostPairing group, two-way RM ANOVA with post-hoc Bonferroni test. (K) Normalized fEPSP_EALS_ slopes before and after HFLSEA/Pre/PostPairing. (L) Example fEPSP_EALS_ traces before and after HFLSEA/Pre/PostPairing. Scale bars: upper, 5 ms and 0.2 mV. (M) Individual and average fEPSP_EALS_ slope changes before and after HFLSEA/Pre/PostPairing. paired t-test, t _(12)_ = -1.424, n.s. p = 0.180, n= 13. See Table S1 for detailed Statistics.

The VC→AC inputs were potentiated when we applied HFLS on the EC→AC CCK^+^ projection but not on the VC→AC projection (Figures 2E, LTP curves; 2F, fEPSP traces; 2G, two-way RM ANOVA, F _(1,21)_ = 10.490, significant interaction, p = 0.004; red, HFLSEA/Pre/PostPairing before [99.9±1.5%] vs. after [120.7±4.0%], pairwise comparison, p < 0.001, n = 13; green, HFLSVA/Pre/PostPairing before [99.9±2.0%] vs. after [97.2±6.7%], pairwise comparison, p = 0.623, n = 10; HFLSEA/Pre/PostPairing after vs. HFLSVA/Pre/PostPairing after, pairwise comparison, p = 0.005). However, simple pairing of HFLS of EC→AC, CCK^+^ projection with single pulse VALS did not potentiate the VC→AC input (Figure S2D, t _(8)_ = -0.899, p = 0.395, n = 9, paired t-test). This may be interpreted as lack of Hebbian postsynaptic activation and agree with our previous study (Li et al., 2014). Besides, fEPSPs evoked by noise stimuli (acoustic stimuli, AS) were also potentiated after HFLSEA/Pre/PostPairing but not HFLSVA/Pre/PostPairing (Figures 2H, LTP curves; 2I, fEPSP traces; 2J, two-way RM ANOVA, F _(1,16)_ = 9.711, significant interaction, p = 0.007; black, HFLSEA/Pre/PostPairing before [100.4±1.2%] vs. after [143.2±14.9%], pairwise comparison, p = 0.001, n = 8; gray, HFLSVA/Pre/PostPairing before [101.7±0.5%] vs. after [100.5±2.3%], pairwise comparison, p = 0.898, n = 10; HFLSEA/Pre/PostPairing after vs. HFLSVA/Pre/PostPairing after, pairwise comparison, p = 0.006). However, the EC→AC, CCK^+^ inputs were not significantly potentiated after HFLSEA/Pre/PostPairing (Figures 2K, LTP curves; 2L, fEPSP traces; 2M, paired t-test, t _(12)_ = -1.424, p = 0.180, n = 13). The results suggest that HFLS of EC→AC CCK^+^ projection rather than VC→AC CaMKII^+^ projection is necessary to induce potentiation of VC→AC inputs, whereby CCK release induced by the former one is an underpinning mechanism. Taken together, our results demonstrate a typical form of heterosynaptic plasticity, in which the potentiation of the VC→AC input is not dependent on HFLS of its own pathway but requires HFLS of the EC→AC projection that presumably triggers CCK release.

### HFLS of EC→AC CCK^+^ terminals in vitro leads to LTP of VC→AC inputs after pairing with electrical stimulation

We also performed similar experiments at the single cell level in vitro. Slices were prepared from CCK-iRES-Cre mice after injection of AAV9-Ef1α-Flex-Chronos-GFP in the EC and of AAV9-Syn-ChrimsonR-tdTomato in the VC (Figure 3A). Pyramidal neurons in the auditory cortex were patched (Figure 3B), and excitatory postsynaptic currents evoked by VALS (EPSC_VALS_, Figure 3C) and electrical stimulation of the auditory cortex (EPSCESAC, Figure 3D) were recorded. HFLS of the EC→AC CCK^+^ projection (Figure 3E) was followed by the pairing of VALS and postsynaptic activation evoked by ESAC, which was repeated for 5 times with an ITI of 10s (Figure 3F left). As a control, we replaced the HFLS of EC→AC CCK^+^ projections with HFLS of VC→AC projection (Figure 3F right).

**Figure 3.**
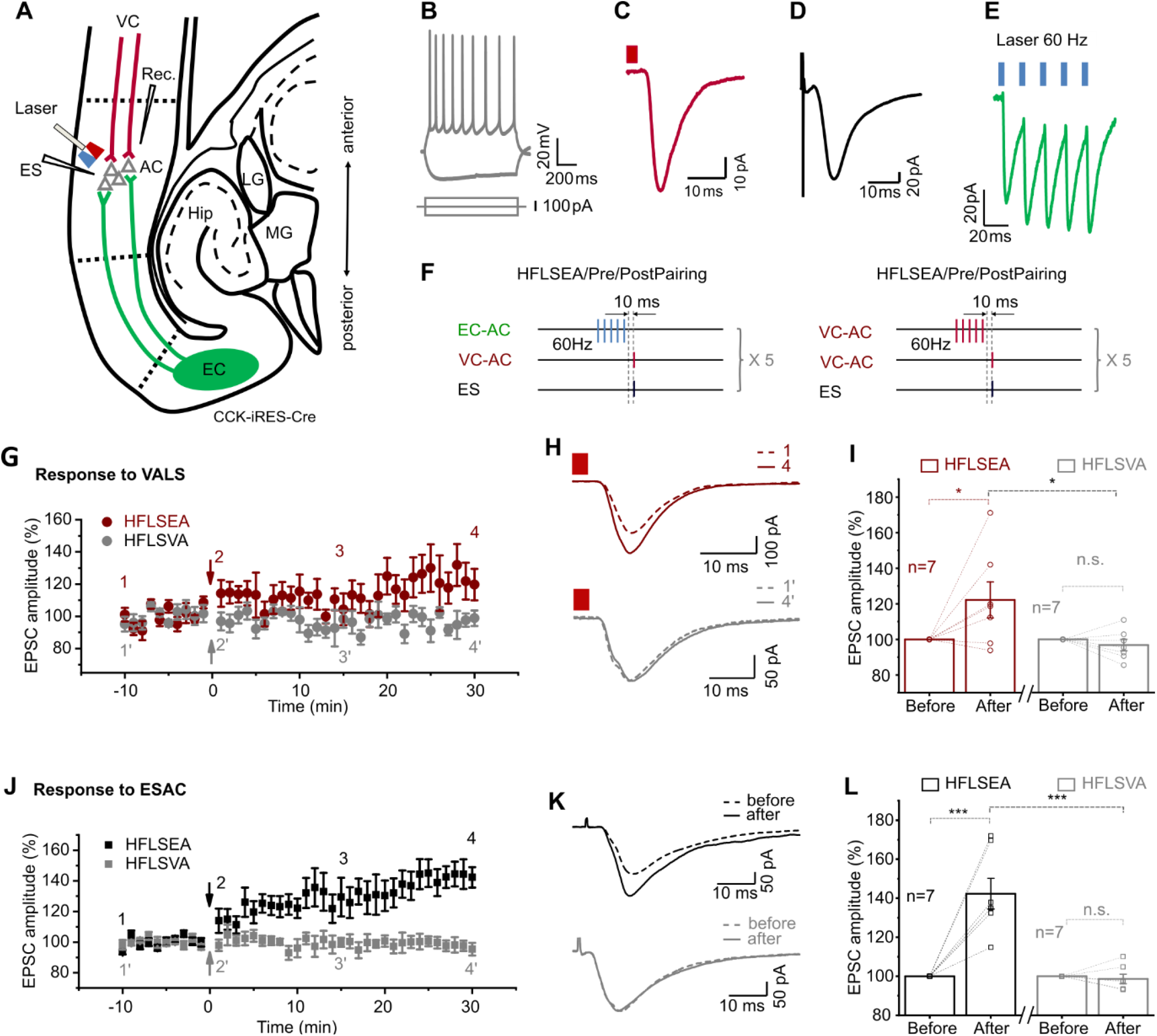
HFLS of ENT→AC CCK^+^ terminals in vitro leads to LTP of VC→AC inputs after pairing with electrical stimulation. (A) Positions of the whole cell recording pipette, electrical stimulation electrode, and the optical fiber in a slice of CCK-iRES-Cre mice with AAV-Ef1α-Flex-Chronos-GFP injected in Ent and AAV-Syn-ChrimsonR-tdTomato injected in VC. (B) Representative pyramidal neuron firing in response to current injection. (C and D) Representative traces of EPSC_VALS_ (C) and EPSC_ESAC_ (D) of pyramidal neuron. Scale bars: C, 10 ms and 10 pA; D, 10 ms and 20 pA. (E) Representative EPSC trace in response to HFLS of Ent→AC CCK^+^ terminals (blue rectangles, 60 Hz, 3 ms/pulse, 5 pulses). (F) Protocols of HFLSEA/Pre/PostPairing (left) and HFLSVA/Pre/PostPairing (right). (G) Normalized EPSC_VALS_ amplitudes before and after HFLSEA/Pre/PostPairing (red) or HFLSVA/Pre/PostPairing (gray). (H) Example EPSC_VALS_ traces before (dashed, at timepoint 1 or 1’ in G) and after (solid, at timepoint 4 or 4’ in G) HFLSEA/Pre/PostPairing (red, upper) or HFLSVA/Pre/PostPairing (gray, bottom). (I) Individual and average EPSC_VALS_ amplitude changes before and after HFLSEA/Pre/PostPairing (red) or HFLSVA/Pre/PostPairing (gray). * p < 0.05, n.s. p = 0.670, n = 7 for HFLSEA/Pre/PostPairing group, n = 7 for HFLSVA/Pre/PostPairing group, two-way RM ANOVA with post-hoc Bonferroni test. (J) Normalized EPSC_ESAC_ amplitudes before and after HFLSEA/Pre/PostPairing (black) or HFLSVA/Pre/PostPairing (gray). (K) Example EPSC_ESAC_ traces before (dashed, at timepoint 1 or 1’ in J) and after (solid, at timepoint 4 or 4’ in J) HFLSEA/Pre/PostPairing (black, upper) or HFLSVA/Pre/PostPairing (gray, bottom). (L) Individual and average EPSC_ESAC_ amplitude changes before and after HFLSEA/Pre/PostPairing (red) or HFLSVA/Pre/PostPairing (gray). *** p < 0.01, n.s. p = 0.814, n = 7 for HFLSEA/Pre/PostPairing group, n = 7 for HFLSVA/Pre/PostPairing group, two-way RM ANOVA with post-hoc Bonferroni test. See Table S1 for detailed Statistics.

Similar to the *in vivo* results, the amplitude of EPSCVALS significantly increased after HFLSEA/Pre/PostPairing but not after HFLSVA/Pre/PostPairing (Figures 3G, LTP curves; 3H, EPSCVALS traces; 3I, two-way RM ANOVA, F(1,12)= 5.759, significant interaction, p = 0.034; red, increased by 22.2 ± 10.1% after HFLSEA/Pre/PostPairing, pairwise comparison, p = 0.012, n = 7; gray, changed by 3.3 ± 3.2% after HFLSVA/Pre/PostPairing, pairwise comparison, p = 0.670, n = 7; HFLSEA/Pre/PostPairing after [122.2 ± 10.1%] vs. HFLSVA/Pre/PostPairing after [96.7 ± 3.2%], pairwise comparison, p = 0.034; S3A and S3B, 10 successive example traces and their averaged trace of the EPSCs at different time points as shown in Figure 3G). Likewise, the EPSCESAC amplitude increased by 42.4 ± 7.9% after HFLSEA/Pre/PostPairing but not after HFLSVA/Pre/PostPairing (Figures 3J, LTP curves; 3K, EPSCESAC traces; 3L, two-way RM ANOVA, F(1,12)=28.074, significant interaction, p < 0.001; black, HFLSEA/Pre/PostPairing before vs. after, pairwise comparison, p < 0.001, n = 7; gray, changed by 1.4 ± 5.8% after HFLSVA/Pre/PostPairing, pairwise comparison, p = 0.814, n = 7; HFLSEA/Pre/PostPairing after [142.4 ± 7.9%] vs. HFLSVA/Pre/PostPairing after [98.6 ± 2.5%], pairwise comparison, p < 0.001; S3C and S3D, 10 successive traces and their averaged trace at different timepoints as shown in Figure 3J). These results, from recording at the synaptic level, provide further evidence for the view that HFLS of EC→AC, CCK^+^ projection is a prerequisite to potentiate the VC inputs to the AC.

### EC→AC projecting neurons have higher CCK expression levels than VC→AC projecting neurons

A variety of neuropeptides, including CCK, are expressed in the neocortex, mostly in interneurons (Somogyi and Klausberger, 2005), although CCK is also expressed in projection neurons. The results in the present study indicate that HFLS of the EC→AC, but not of the VC→AC, projection can produce LTP in AC. We next explored if levels of CCK transcript, and thus possibly CCK peptide, could underly this difference by using RNAscope combined with retrograde tracing with AAV virus. We injected AAVretro-hSyn-Cre-WPRE-hGH in the AC of Ai14 mice, a Cre reporter line, retrogradely labeling the EC and VC neurons projecting to AC with Cre-dependent tdTomato. The expression level of CCK was then assessed by RNAscope, a semi-quantitative *in situ* hybridization method (Figure 4A-D). We found that the *expression level* of CCK was significantly higher among projecting neurons in the EC than in the VC across three animals analyzed (Figures 4E and S4). The *proportion* of projecting neurons expressing elevated CCK levels was also higher in the EC compared with the VC (Figure 4F). These results suggest that after HFLS more CCK is released from EC→AC neurons than from VC→AC neurons, which may, at least, be one explanation why the former but not the latter can produce LTP.

**Figure 4.**
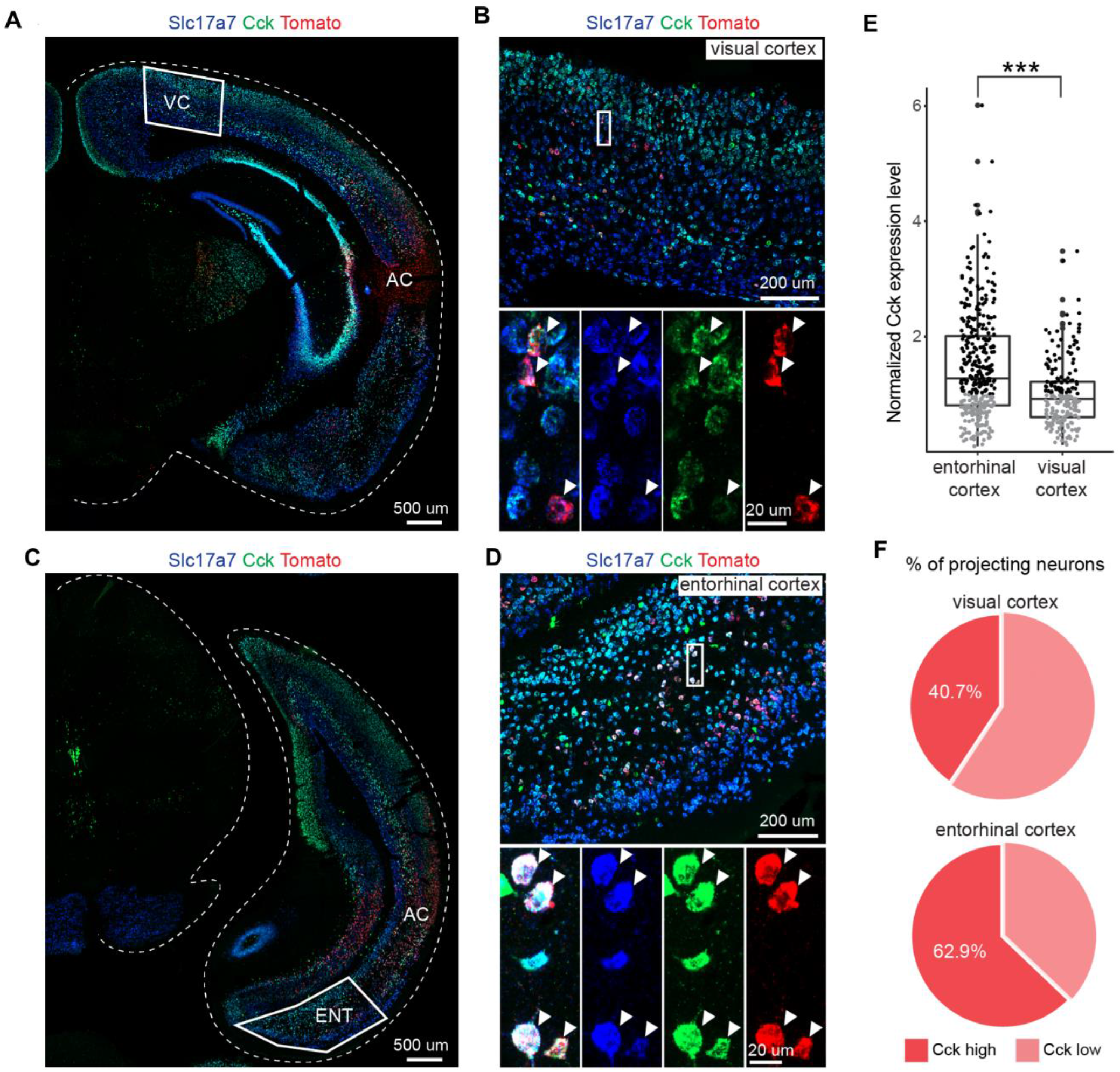
AC projection neurons in the Ent express higher level of CCK than that in the VC. (A) Overview of injection site at the AC and projecting neurons in the VC. Scale bar: 500 um. (B) Expression of Slc17a7 (vGlut1) and Cck in retrogradely labeled neurons (tdTomato+) in the VC. Scale bars: upper, 200 μm; bottom, 20 μm. (C) Overview of injection site at the AC and projecting neurons in the Ent. Scale bar: 500 μm. (D) Expression of Slc17a7 and Cck in retrogradely labeled neurons (tdTomato+) in the EC. Scale bars: upper, 200 μm; bottom, 20 μm. (D) Comparison of Cck expression level in AC projecting neurons of EC and VC (data points are from 3 animals). unpaired t-test, ***p<0.001. Black, high level; gray, low level. (E) Pie chart showing percentage of projecting neurons expressing low and high Cck level in the VC (upper) and the EC (bottom), respectively. See Table S1 for detailed Statistics.

### Effect of different parameters of the pairing protocol on the potentiation level of VC→AC inputs

Neuropeptide release likely is frequency-dependent (Bean and Roth, 1991; Hökfelt, 1991; Iverfeldt et al., 1989; Lundberg and Hokfelt, 1983; Shakiryanova et al., 2005; Whim and Lloyd, 1989), and our results suggest that CCK released from the EC→AC CCK^+^ projection was critical for generating visuo-auditory cortical LTP. We hypothesized that the frequency of the laser used to stimulate the EC→AC CCK^+^ projection was critical for the level of potentiation of the VC→AC input. We therefore varied the *frequency of the laser stimulation* (80, 40, 10, or 1 Hz). As shown in Figure 5A left, the delay between the termination of repetitive laser stimulation of the CCK^+^ EC→AC projection and presynaptic activation (Delay 1) was set at 10 ms, and the delay between pre and postsynaptic activation (Delay 2) was set at 0 ms. The potentiation level of the VC→AC inputs showed a tendency to increase as the frequency of laser stimulation of the CCK^+^ EC→AC projection increased (Figure 5A right, two-way RM ANOVA, F(3,34) = 10.666, significant interaction, p < 0.001; 1 Hz before [99.7 ± 1.3%] vs. after [96.6 ± 2.9%], pairwise comparison, p = 0.352, n =9; 10 Hz before [98.5 ± 1.2%] vs. after [105.8 ± 2.5%], pairwise comparison, p =0.044, n = 8; 40 Hz before [99.4 ± 0.8%] vs. after [110.5 ± 1.8%], pairwise comparison, p = 0.003, n=8; 80 Hz before [99.9 ± 1.5%] vs. after [120.7 ± 4.0%], pairwise comparison, p < 0.001, n = 13). If higher than 10 Hz, the VC→AC input was significantly potentiated. However, at 1 Hz no significant potentiation was observed.

**Figure 5.**
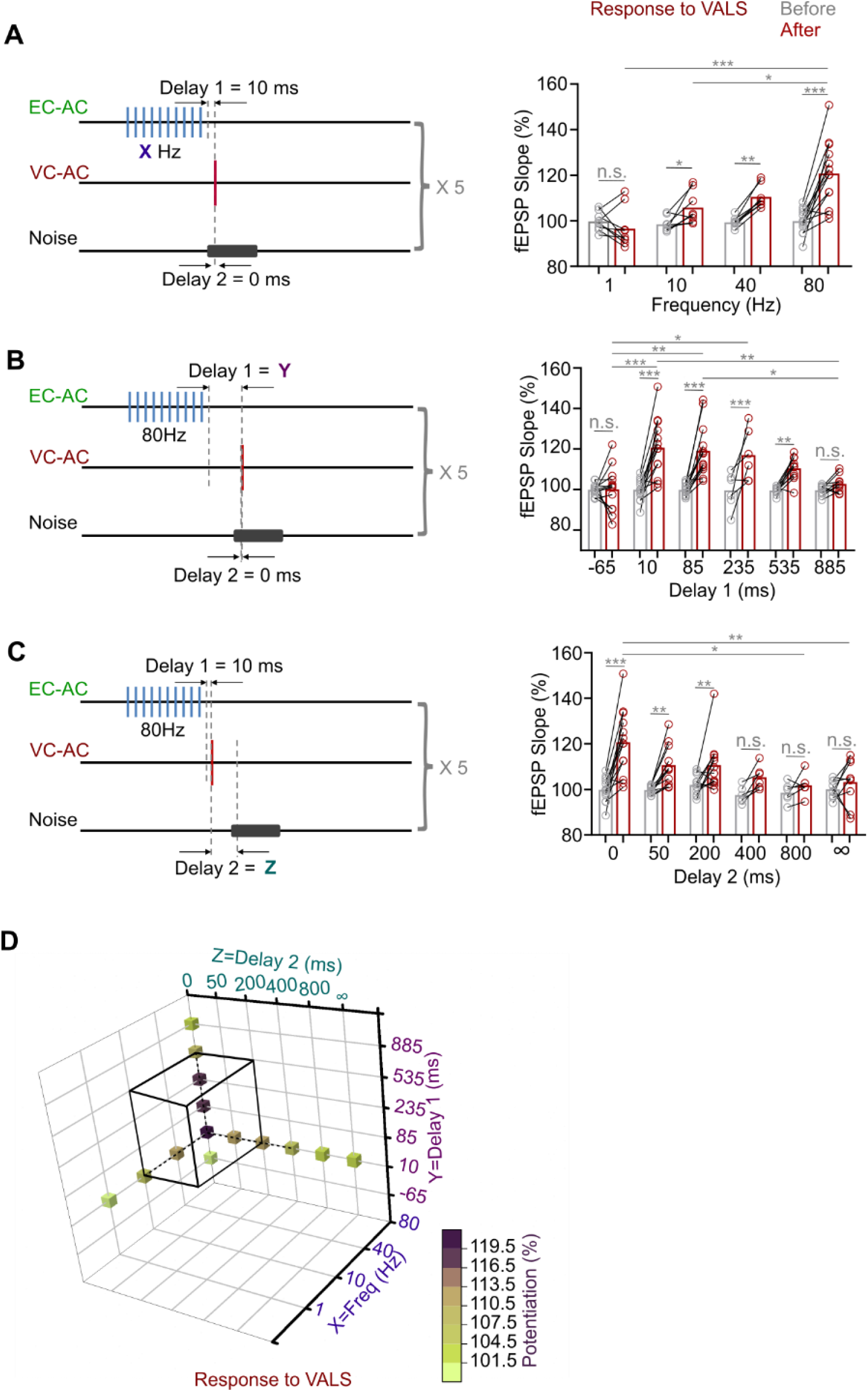
Effect of different parameters of the pairing protocol on the potentiation level of VC→AC inputs. (A) Left: schematic drawing of experiment design. Delay1 =10 ms, Delay2 = 0 ms, and the frequency was varied (80, 40, 10, or 1 Hz). Right: individual and average fEPSP_VALS_ slopes (normalized to the baseline) after pairing at different frequencies. Two-way RM ANOVA with post-hoc Bonferroni test, n.s. no significant, * p < 0.05, ** p < 0.01, *** p < 0.001, n = 9 for 1 Hz, n = 8 for 10 Hz, n = 8 for 40 Hz, n = 13 for 80 Hz. Data points in groups 1 Hz and 40 Hz refer to our previous study (Zhang et al., 2020). (B) Left: schematic drawing of experiment design. HFLS Frequency = 80 Hz, Delay2 = 0 ms, and Delay 1 was varied (10, 85 235, 535, 885, and -65 ms). Right: individual and average fEPSP_VALS_ slopes (normalized to the baseline) after pairing at different Delay1s. Two-way RM ANOVA with post-hoc Bonferroni test, n.s. no significant, * p < 0.05, ** p < 0.01, *** p < 0.001, n=13, 13, 13, 6, 10, 10 for Delay 1=-65, 10, 85, 235, 535, and 885 ms, respectively. (C) Left: schematic drawing of experiment design. HFLS Frequency = 80 Hz, Delay1 = 10 ms, and Delay 2 was varied (0, 50, 200, 400, 800 ms, and ∞). Right: individual and average fEPSP_VALS_ slopes (normalized to the baseline) after pairing at different Delay2s. Two-way RM ANOVA with post-hoc Bonferroni test, n.s. not significant, * p < 0.05, ** p < 0.01, *** p < 0.001, n = 13, 11, 12, 6, 6, 9 for Delay 2= 0, 50, 200, 400, 800 ms and ∞, respectively. (D) Three-dimensional summary of the effect of different parameters (Frequency, Delay1 and Delay2) on the potentiation level of VC→AC inputs. Parameters locate inside black cubes can induce significant potentiation. See Table S1 for detailed Statistics.

In contrast to small-molecule neurotransmitters that are rapidly cleared by reuptake pumps, neuropeptides are mostly released extra-synaptically, are removed/inactivated more slowly, and may have longer-lasting effects. Thus, we explored the role of Delay 1, i.e. if the *time* interval between the termination of HFLS and the Pre/Post Pairing influenced the degree of potentiation of the VC→AC inputs (Delay 2 = 0 ms, HFLS frequency = 80 Hz, Figure 5B left). The VC→AC inputs were significantly potentiated, when Delay 1 was 10, 85, 235, or 535 ms rather than 885 or -65 ms (Figure 5B right, two-way RM ANOVA, F(5,59) =7.115, significant interaction, p < 0.001; 10 ms before [99.9 ± 1.5%] vs. after [120.7 ± 4.0%]], pairwise comparison, p < 0.001, n = 13; 85 ms before [99.8 ± 0.8%] vs. after [119.1 ± 3.5%], pairwise comparison, p < 0.001, n= 13; 235 ms before [99.5 ± 3.7%] vs. after [117.0 ± 5.2%], pairwise comparison, p < 0.001, n= 6; 535 ms before [99.4 ± 0.6%] vs. after [110.4 ± 1.9%], pairwise comparison, p = 0.003, n= 10; 885 ms before [99.6 ± 0.8%] vs. after [102.7 ± 1.3%], pairwise comparison, p = 0.385, n= 10; -65 ms before [99.0 ± 0.8%] vs. after [100.1 ± 3.0%], pairwise comparison, p = 0.945, n= 13).

The Hebbian theory states, popularly, that “cells that fire together wire together”(Löwel and Singer, 1992), a more accurate interpretation being ‘synaptic strength increases when the presynaptic neuron always fires immediately before the post-synaptic neuron’(Caporale and Dan, 2008). Based on this, the interval between pre and postsynaptic activation should be critical for potentiation. In the next experiment, the interval (Delay 2) between VC→AC projection activation (i.e., presynaptic activation) and natural auditory cortex activation (i.e., postsynaptic activation) was set as the only variable (Delay 1 = 10 ms, HFLS frequency = 80 Hz, Figure 5C left). The potentiation of VC→AC inputs showed a trend towards decrease as Delay 2 increased. Significant potentiation was observed when Delay 2 was 0, 50, 200, rather than 400, 800 ms and ∞ (without noise) (Figure 5C right, two-way RM ANOVA, F(5,51) =4.133, significant interaction, p = 0.003; 0 ms before [99.9 ± 1.5%] vs. after [120.7 ± 4.0%]], pairwise comparison, p < 0.001, n = 13; 50 ms before [99.7 ± 0.6%] vs. after [110.7 ± 2.7%], pairwise comparison, p = 0.001, n= 11; 200 ms before [102.0 ± 1.2%] vs. after [110.6 ± 3.3%], pairwise comparison, p = 0.006, n= 12; 400 ms before [97.5 ± 1.5%] vs. after [105.3 ± 2.1%], pairwise comparison, p =0.073, n= 6; 800 ms before [98.6 ± 1.8%] vs. after [101.8 ± 2.1%], pairwise comparison, p = 0.454, n= 6; ∞ before [100.0 ± 1.3%] vs. after [103.2 ± 3.3%], pairwise comparison, p = 0.363, n= 9).

Taken together, to induce a significant potentiation of VC→AC inputs (Figure 5D, black cube) within five trails pairing with an ITI of 10 s, (i) the frequency of repetitive laser stimulation of the CCK^+^ EC→AC projection should be 10 Hz or higher, (ii) Delay 1 should be longer than 10 ms and shorter than 535 ms, and (iii) Delay 2 should be longer than 0 ms and shorter than 200 ms.

### Application of a CCKBR antagonist blocks generation of the visuo-auditory association

The above results showed that the generation of visuo-auditory associations needed inputs from the EC together with high-frequency stimulation, presumably triggering CCK release from their terminals. Next, we ask whether or not CCK is essential for generation of visuo-auditory associations, which can be reflected in a behavioural context.

To that end, we examined the role of CCK in the formation of visuo-auditory associative memory in a fear response test (Figure 6A). First, a CCKB receptor (CCKBR) antagonist (L-365,260) or ACSF was injected bilaterally into the AC, followed by a 25-trial pairing session of the visual stimulus (VS) with the AS. We repeated the above drug and pairing session 4 times per day and on 3 consecutive days. On day 4, baseline tests for the freezing response to the AS and VS were performed (3 trials) before the mouse was fear conditioned to the AS. After fear conditioning, freezing responses to the AS and VS were further examined on day 5 (Figure 6A). As expected, mice showed no freezing response to the AS before conditioning, but a high freezing rate to the AS after conditioning (Video S3, S4, S7, S8; Figure 6B, ACSF-AS-Baseline [7.4 ± 3.2%] vs. ACSF-AS-Post intervention [67.8 ± 4.0%], p < 0.001, n = 9; L-365,260-AS-Baseline [5.8 ± 1.6%] vs. L-365,260-AS-Post intervention [67.5 ± 4.1%], p < 0.001, n = 8; two-way RM ANOVA with post-hoc Bonferroni test). The ACSF mice group showed a significantly increased freezing response to the VS, indicating that an association between the AS and VS had been established during the pairings (Videos S1, S2; Figure 6B blue, ACSF-VS-Baseline [4.1 ± 0.9%] vs. ACSF-VS-Post intervention [24.4 ± 3.8%], p<0.001, n = 9, two-way RM ANOVA with post-hoc Bonferroni test). However, the bilateral infusion of L-365,260 into the AC blocked this association, resulting in a nil response to the VS (Video S5, S6; Figure 6B red, L-365,260-VS-Baseline [4.3 ± 1.2%] vs. L-365,260-VS-Post intervention [2.5 ± 1.6%], p = 1.000, n = 8, two-way RM ANOVA with post-hoc Bonferroni test). Mice showed a promising association between the AS and foot shock, as indicated by high freezing rate following the AS. However, an association between the VS and AS was not established. There was a significant difference between the freezing rates to the VS of experimental and control groups (Figure 6B; L-365,260-VS-Post intervention [2.5 ± 1.6%, n = 8] vs. ACSF-VS-Post intervention [24.4 ± 3.8%, n = 9], p < 0.001, two-way RM ANOVA with post-hoc Bonferroni test). These results demonstrate that the CCKBR antagonist prevented the generation of the association between the VS and AS and suggest an essential role of CCK in the generation of the visuo-auditory association.

**Figure 6.**
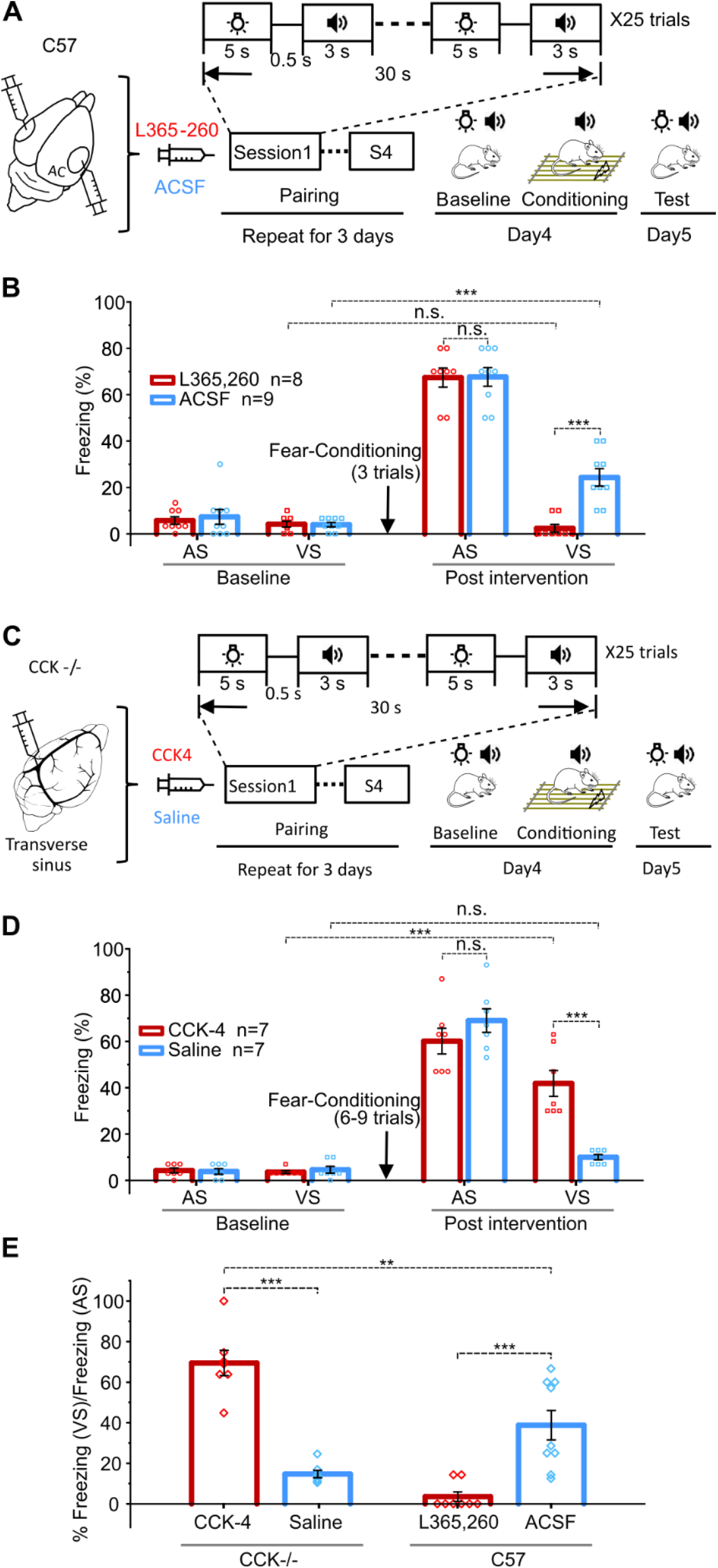
Visuo-auditory associative memory could not be formed without CCK. (A) Schematic drawing of the training protocol for wild-type mice to associate the VS and AS. L-365,260 or ACSF was infused to the AC before pairing of VS and AS. (B) Bar chart showing freezing percentages to the AS and VS before and after the conditioning. *** p<0.01, n.s. p = 1.000 for L-365,260-VS-Baseline vs. L-365,260-VS-Post intervention, n.s. p = 0.962 for L-365,260-AS-Post intervention vs. ACSF-AS-Post intervention, n=8 for L-365,260 group, and n = 9 for ACSF group, two-way RM ANOVA with post-hoc Bonferroni test. (C) Schematic drawing of the training protocol for CCK^-/-^ mice to associate the VS and AS. CCK-4 or ACSF was systemically administrated through the drug infusion cannula in the transverse sinus before paring of VS and AS. (D) Bar chart showing freezing percentages of CCK^-/-^ mice to the VS and AS before and after the intervention. *** p<0.01, n.s. p = 1.000 for Saline-VS-Baseline vs. Saline-VS-Post intervention, n.s. p = 0.264 for CCK-AS-Post intervention vs. Saline-AS-Post intervention, n=7 for CCK-4 group, and n = 7 for Saline group, two-way RM ANOVA with post-hoc Bonferroni test. (E) The ratio of freezing in response to the VS over that to the AS after the intervention of wild-type and CCK-/- in all conditions are summarized in the bar chart. ** p<0.01, *** p<0.001, one-way ANOVA with post-hoc Bonferroni test. See Table S1 for detailed Statistics.

### Systemic administration of CCK4 rescues the deficit of CCK-/- mice in visuo-auditory memory

As seen after treatment with a CCK antagonist, we expected that CCK^-/-^ mice would show a deficit in the formation of associative memory, and we tested this hypothesis (Figure 6C). Our previous results demonstrated that 6-9 trials were needed for CCK^-/-^ mice to produce a freezing rate of >60% in response to the conditioned AS, whereas only 3 trials were needed for wild-type mice, suggesting a general associative learning deficit in the CCK^-/-^ mice (Chen et al., 2019). The CCK^-/-^ mice in the control group (with saline injection) consistently showed a minimal freezing response to VS after visuo-auditory association and fear conditioning (Videos S9-12; Figure 6D blue). To determine whether systemic administration of CCK could rescue this deficit, we administrated CCK-4 through a drug infusion cannula implanted into the transverse sinus. There is evidence that the tetrapeptide CCK-4 can penetrate the blood-brain barrier (Rehfeld, 2000). The CCK-4 dosage (1ug/kg) was at sub-panic attack level, and animals showed no sign of panic or anxiety after administration (Figure S5). CCK-4 injection resulted in a significantly higher freezing rate compared to the controls (Videos S13-16; Figure 6D red, CCK-VS-Post intervention [41.9 ± 5.6%, n = 7] vs. Saline-VS-Post intervention [10.0 ± 1.1%, n = 7], p < 0.001; two-way RM ANOVA with post-hoc Bonferroni test), indicating that the visuo-auditory association was rescued upon CCK4 administration.

To better compare the strength of visuo-auditory association under different experimental conditions, we calculated the ratio of the freezing response to the VS compared with that to the AS after conditioning (Figure 6E). The ratio of the CCKBR antagonist (L-365,260)-treated group was lowest among all groups, demonstrating a nearly complete abolishment of the visuo-auditory association. Interestingly, the ratio of the CCK-4 group was the highest among all groups (Figure 6E. F (3, 27) = 28.797, CCK-4 infusion in CCK^-/-^ mice [69.5 ± 6.2%, n = 7] vs. ACSF infusion in the wild-type mice [38.8 ± 7.3%, n = 9], p = 0.002, one-way ANOVA with post-hoc Bonferroni test). This result indicates a possible compensatory upregulation of CCK receptors in CCK^-/-^ mice, leading to the highest association between the VS and AS, findings that are worth further investigation.

## Discussion

In the present study we demonstrate, in the mouse, that a direct input from the VC to the AC can be potentiated after pairing with postsynaptic firing evoked by auditory stimuli in the presence of CCK. This was observed both after exogenous administration and presumably after endogenous CCK release after HFLS of the CCK^+^ EC→AC projection. In fact, a significant potentiation of the presynaptic input could be induced after only five trials of pairing, even if it arrived 200 ms earlier than the postsynaptic activation. Behavioral experiments proved that a CCKBR antagonist could block the generation of the visuo-auditory association, whereas a CCK agonist could rescue the defect association seen in CCK^-/-^ mice.

### Critical projections

Cross-modal association can be considered as the potentiation of synaptic strength between different modalities. Consistent with other studies(Bizley et al., 2007; Budinger et al., 2006; Falchier et al., 2002; Falchier et al., 2010; Rockland and Ojima, 2003), we describe a direct projection in the mouse from the VC to the AC for the visuo-auditory association using both retrograde and anterograde tracing methods. The projection terminates both in the superficial and deep cortical layers. Our previous study also on mouse demonstrated that neurons in the entorhinal cortex retrogradely labeled by true blue injected into the AC are almost 100% CCK^+^ (Li et al., 2014). We here confirm that CCK also is expressed in neurons of the EC projection to the AC.

### Cortical neuropeptides

Cortical neurons express a number of neuropeptides (Somogyi and Klausberger, 2005), whereby CCK is the most abundant of all. CCK comes in different forms, but it is the sulphated octapeptide, CCK-8S, that predominates in the brain (Dockray et al., 1978; Rehfeld, 1978). CCK is expressed in GABAergic interneurons (Houser et al., 1983), and many pyramidal neurons (DeFelipe and Fariñas, 1992) have also high levels of CCK transcript (Burgunder and Young, 1988; Schiffmann and Vanderhaeghen, 1991). The CCK^+^ interneurons are relatively few, but exert a critical control of cortical activity (Somogyi and Klausberger, 2005). However, it is CCK in the pyramidal neurons that are in focus in the present study, especially the CCK^+^, glutamatergic projection from EC to AC. We also use exogenous CCK in the experiments, both CCK-8S and CCK-4 which is the C-terminus not only of proCCK but also of gastrin. The small size of the latter fragment is the reason, why it is considered to pass the blood-brain-barrier (Rehfeld, 2000). However, we infused CCK-4 into the transverse venous sinus aiming at obtaining maximal peptide levels in the cortex.

### Optogenetics

The present study is based on optogenetics, that is genetic introduction of light sensitive channels (Channelrodopsins), allowing control of selective neuron populations by light - a method that has revolutionized neuroscience research(Deisseroth et al., 2006; Knopfel et al., 2010). Here, using the two channels Chronos and ChrimsonR, we were able to activate two distinct projection terminals converging in the same target area activate (the EC→AC CCK^+^ projection and the VC→AC projection).

### Prerequisits for synaptic plasticity

Our previous finding based on in vivo intracellular recording indicated that there are three prerequisites to enable synaptic plasticity: presynaptic activation, postsynaptic firing, and, in this particular system, the presence of CCK(Li et al., 2014). Replacing the classical HFS protocol (HFLS of VC→AC projections) with local infusion of CCK-8S followed by pairing between pre-synaptic VC→AC inputs induced by VALS and postsynaptic firing evoked by noise stimuli led to the potentiation of VC→AC inputs. We hypothesize that these events may underlie the visuo-auditory association observed in the auditory cortex, further demonstrating the critical role of CCK to enable synaptic plasticity(Chen et al., 2019; Li et al., 2014).

Simple pairing between HFLS of EC→AC, CCK^+^ projection and VALS without postsynaptic activation did not induce LTP in VC→AC inputs. Neither did low frequency (1 Hz) laser stimulation of the EC→AC CCK^+^ projection induce LTP of VC→AC inputs, probably since no/not enough CCK was released by low frequency stimulation. Surprisingly, no LTP was recorded after HFLS of the VC→AC projection followed by Pre/Post Pairing. We demonstrate that VC→AC projecting neurons have relatively lower CCK expression compared to the EC→AC projecting neurons. This could be a reason why LTP was not observed for the VC→AC inputs after only 5 trails of HFLSVA/Pre/PostPairing. If the number of pairing trials or if laser stimulation intensity reaches a certain level, enough CCK may be released from the CCK^+^ VC→AC projecting neurons, and LTP may occur.

These findings suggest that in traditional LTP HF stimulation also activates CCK^+^ projection terminals, thereby releasing CCK and enabling potentiation (Chen et al., 2019). Subsequent experiments, in which the frequency of repetitive laser stimulation of EC→AC CCK^+^ projection terminals was changed, showed that the degree of potentiation increased with increasing frequency. This finding can be explained by the frequency-dependent nature of neuropeptide (CCK) release(Bean and Roth, 1991; Hökfelt, 1991; Iverfeldt et al., 1989; Shakiryanova et al., 2005; Whim and Lloyd, 1989).

In addition, the potentiation of VC→AC inputs decreased as the interval between the termination of HFLS and Pre/Post Pairing increased in the positive direction. A time window of 535 ms was observed to produce significant potentiation. If we would have increased the number of pairing trials, the effective Delay 1 might have lengthened.

### Hebbian plasticity

We addressed the issue of spike timing-dependent plasticity and the critical time limit. Here the Hebbian rule has, arguably, been the most influential theory in learning and memory. This rule says that in order to induce potentiation the presynaptic subthreshold input should occur at most 20 ms before post-synaptic firing (Bi and Poo, 1998; Markram et al., 1997; Zhang et al., 1998). However, this theory has been challenged (Drew and Abbott, 2006; Izhikevich, 2007), with a prevalent question being: how can associations be established across behavioral time scales of seconds or even longer if the critical window is only 20 ms? (Bittner et al., 2017). Bittner et al. reported that five pairings of pre-synaptic subthreshold inputs with post-synaptic calcium plateau potentials produce a large potentiation; and that presynaptic inputs can arrive seconds before or after postsynaptic activity, a phenomenon termed “behavioral time scale synaptic plasticity” (Bittner et al., 2017).

This may account for the highly plastic nature of place fields in the hippocampus. In agreement, we observed potentiation in the neocortex, even when presynaptic input arrived 200 ms (Delay 2) earlier than postsynaptic firing and after only five trials pairing, but only in the presence of CCK. This time window could perhaps be extended as the paring trials increase. In general terms, these results fit with the fact that neuropeptides are known to exert slow and long-lasting effects (van den Pol, 2012). We did not explore the reverse direction, in which post-synaptic activity occurred before pre-synaptic activity, because noise stimuli induced more than one spike and thus timing would be difficult to control.

In summary, we found that a direct projection from the VC to the AC provide an anatomical basis for visuo-auditory association. The VC→AC inputs was potentiated after pairing with postsynaptic firing evoked by the auditory stimulus in the presence of CCK that was applied either exogenously or (endogenously) released from the EC→AC CCK^+^ projection terminals stimulated with a HF laser. In the presence of endogenous CCK, significant potentiation of presynaptic input could be induced even it arrived 200 ms earlier than postsynaptic firing and after only five trials of pairing. Finally, through the behavior experiments, we proved that the CCKBR antagonist blocked the establishment of visuo-auditory association, whereas the CCK agonist rescued the deficit of association in the CCK^-/-^ mouse.

## Materials and Methods

### Animals

Adult male and female Sprague Dawley rats, C57BL/6J, CaMKIIa-Cre (Jackson lab stock #005359), Ai14 (Jackson lab stock #007914), and *CCK-ires-Cre* (Jackson lab stock #012706) mice were used in current study. For behavior experiments, only male animals were used. Animals were confirmed to have clean external ears and normal hearing and were housed in a 12-hour-light/12-hour-dark cycle (lights on from 8:00 pm to 8:00 am next day) at 20-24°C with 40-60% humidity with ad libitum access to food and water. All procedures were approved by the Animal Subjects Ethics Sub-Committees of City University of Hong Kong.

### Anesthesia and surgery

To induce anesthesia for retrograde tracer or virus injection, animals were administered (i.p.) with 50 mg/kg pentobarbital (Dorminal 20%, Alfasan International B.V., Woerden, Netherlands). To induce anesthesia for acute *in vivo* recording, urethane sodium (2 g/kg, i.p., Sigma-Aldrich, St. Louis, MO, USA) was used with periodic supplements throughout surgery and neuronal recordings. During surgery, lidocaine (2%, Tokyo Chemical Industry [TCI] #L0156, Tokyo, Japan) was frequently applied in drops on the incision site. After confirming anesthesia, head fur between the eyes and ears was trimmed. Animals were mounted on a stereotaxic instrument (for rats, Narishige # SR-6R-HT, Japan; for mice, RWD Life Science # 68001, China). The scalp was sterilized with 70% ethanol. The body temperature was maintained at 37 – 38°C with a heating blanket (Homeothermic Blanket system, Harvard Apparatus, US) during surgery. After making a midline incision, muscle or periosteum was carefully removed with a scalpel blade. And after leveling, we made craniotomies over different target brain regions. All surgery tools were autoclaved before experiments. Before returning to the Laboratory Animal Research for normal holding, we monitored animals until they completely regained consciousness. To prevent infection, we applied erythromycin ointment to the wound for 7 days after surgery.

### Retrograde tracing

For retrograde tracing on rat, a craniotomy window of 1.5 mm by 2.5 mm was made to access the auditory cortex at the left temporal bone, 3.0 to 4.5 mm from the top edge (dorsal-ventral, DV) and −3.0 to −5.5 mm from bregma (anterior-posterior, AP). CTB 488 (1 mg/mL, Molecular Probes #C34775, US) was injected into three locations with different coordinates (AP -3.5 mm, DV -3.8 mm; AP -4.5 mm, DV -3.8 mm; and AP-5.5, DV -3.8). For each location, two depths from the surface of the cortex were adopted (−500 μm and -900 μm, 50 nl each depth). After all injections, the craniotomy window was filled with Kwik-cast silicone gel (World Precision Instruments, US), and the incision was sutured. Animals were kept on the heating pad until voluntary movements, and then returned to their cages. Five days later, animals were transcardially perfused with PBS and 4% paraformaldehyde, sequentially.

### in vivo fEPSP Recording with optogenetics

For dual pathways, we injected AAV9-Ef1α-Flex-Chronos-GFP (3.7 E+12 vg/mL, Boyden/UNC vector core) and AAV9-Syn-ChrimsonR-tdTomato (4.1 E+12 vg/mL, Boyden/UNC vector core) in the Ent and VC of CCK-ires-Cre mic*e* to separately activate auditory cortical projections from entorhinal and visual cortices respectively. We tried both the lateral (LENT, AP -4.2 mm, ML 3.5 mm and DV -3.0 mm [below the pia], 300 nl) and medial (MENT, AP -4.9 mm, ML 3.3 mm, DV -3.2, -2.5 and -1.8 mm [below the pia, 7°to the rostral direction], 100 nl each depth) part of Ent.. For the VC, to avoid the spread of virus directly into the auditory cortex, the medial part of the visual cortex was chosen as the target area. Two locations distributed rostro-caudally with different coordinates (AP -2.7 mm, ML 1.7 mm, DV -0.5 mm; AP -3.3 mm, ML 1.7 mm, DV - 0.5 mm; 150 nl each depth) were adopted. Before the electrophysiological experiment, we waited for 4-5 weeks for the opsins to well expressed in the projection terminals. We adopted 473 nm and 635 nm lasers to stimulate the projection terminals in the auditory cortex from the entorhinal and visual cortices respectively and recorded the laser evoked fEPSP with glass pipette electrodes (∼1M Ohm) rather than metal electrodes to prevent photoelectric artifacts. During recording, the optic fiber was positioned on the surface of the auditory cortex to illuminate the recording site. To examine the optimal laser intensity needed for separately activating the two channelrhodopsins, we injected AAV9-Ef1α-Flex-Chronos-GFP in the Ent (Figure S2A), or AAV9-hSyn-ChrimsonR-tdTomato in the VC (Figure S2B) of the CCK-iRES-Cre mice. In the former, as shown in Figure S2A right, the fEPSP slopes gradually increased with increasing intensity of the 473 nm laser and became saturated at 30 mW/mm^2^ (green solid). However, no responses were evoked by the 635-nm laser, even at an intensity of 40 mW/mm^2^ (red dash). Conversely, in animal with injection of AAV9-hSyn-ChrimsonR-tdTomato in the VC the fEPSP slopes gradually increased and became saturated (red solid) with the 635 nm laser. However, here 40 mW (green dash) produced fEPSPs, but they were relatively small (Figure S2B right). Thus, to avoid cross talk we controlled the fiber end intensities of 473 nm and 635 nm laser below 30 and 40 mW/mm^2^.

For single pathway (VC→AC) activation experiment, we injected AAV9-Ef1α-Flex-Chronos-GFP and AAV9-Syn-ChrimsonR-tdTomato in the VC of CaMKIIα-Cre and wildtype mice respectively. The coordinates are same as above.

### Brain slice preparation and patch clamp recordings

At least four weeks after virus injection, acute brain slices were prepared using a protective cutting and recovery method to achieve a higher success rate for patch clamp. Briefly, anesthetized mice received transcardial perfusion with NMDG-aCSF (92 mM NMDG, 2.5 mM KCl, 1.25 mM NaH2PO4, 30 mM NaHCO3, 20 mM HEPES, 25 mM glucose, 2 mM thiourea, 5 mM Na-ascorbate, 3 mM Na-pyruvate, 0.5 mM CaCl2·4H2O and 10 mM MgSO4·7H2O; pH 7.3-7.4), and the brain was gently extracted from the skull and then cut into 300 μm thick sections. Slices were submerged in NMDG-aCSF for 5-10 min at 32-34 °C to allow protective recovery and then incubated at room-temperature ACSF (119 mM NaCl, 2.5 mM KCl, 1.25 mM NaH2PO4, 24 mM NaHCO3, 12.5 mM glucose, 2 mM CaCl2·4H2O and 2 mM MgSO4·7H2O, ∼25 °C) for at least 1 h before transferring into recording chamber. All solutions were oxygenated with 95% O_2_/5% CO_2_ for 30 min in advance.

Whole-cell recordings were made from pyramidal neurons at the auditory cortex at room temperature. The signals were amplified with Multiclamp 700B amplifier, digitized with Digital 1440A digitizer and acquired at 20 kHz using Clampex 10.3 (Molecular Devices, Sunnyvale, CA). Patch pipettes with a resistance between 3-5 MΩ were pulled from borosilicate glass (WPI) with a Sutter-87 puller (Sutter). The intracellular solution contained: 145 mM K-Gluconate, 10 mM HEPES, 1 mM EGTA, 2 mM MgATP, 0.3 mM Na2-GTP, and 2 mM MgCl2; pH 7.3; 290– 300 mOsm. The pipette was back-filled with internal solution containing 145 mM K-Gluconate, 10 mM HEPES, 1 mM EGTA, 2 mM MgATP, 0.3 mM Na2-GTP, and 2 mM MgCl2; pH 7.3; 290–300 mOsm.

Pyramidal neurons were selected based on the pyramidal-like shape and the firing pattern of regular spiking by injecting a series of hyperpolarizing and depolarizing current with 50 pA increments (1s). An electrical stimulation electrode was placed ∼200 μm from the recording neuron and laser stimulation were presented by using Aurora-220 (473 nm and 635 nm, NEWDOON) through an optic fiber. Only neurons that could be activated by laser stimulation and electrical stimulation were recruited in the following experiment. Under voltage-clamp recording mode (holding at -70 mV), EPSCs induced by electrical stimulation (0.05 Hz, 0.5 ms) and laser stimulation (0.05 Hz, 1-5 ms, 2mW) were stably recorded for 10 min. Then high frequency (60 Hz, 5 pulses, 1-5 ms duration, 2 mW) stimulation of CCK terminal or VC terminal was delivered, followed by the pairng of simultaneous VALS and ESAC. This stimulation protocol was repeated 5 times with a 10-s interval. Electrical stimulation induced EPSCs were then recorded for another 30 min. Intensities of electrical stimulation were adjusted to maintain the amplitude of EPSCs at around 200-300 pA. -5 mV hyperpolarizing pulses (10 ms) were applied every 20s to measure access resistance (Ra) throughout whole recording procedure and recordings were terminated if Ra changed more than 20%.

### Quantification of CCK expression levels of AC projecting neurons in the VC and the Ent

AAVretro-Cre (pENN-AAV/retro-hSyn-Cre-WPRE-hGH, 2.10E+13 vg/mL, Addgene, USA) was injected in three locations (100 nl/each) of the left AC of the Ai14 mice. The coordinates were as follows: AP -2.6 mm (site 1) or -2.9 mm (site 2) or -3.2 mm (site 3), ML 1.0 mm ventral to the edge differentiating the parietal and temporal skull, and DV -0.5 mm from the dura. Three weeks later, the mice were deeply anesthetized with pentobarbital (Dorminal 20%, Alfasan International B.V., Woerden, Netherlands) and transcardially perfused with 20 ml of warm (37°C) 0.9% saline, 20 ml of warm fixative (4% paraformaldehyde, 0.4% picric acid, 0.1% glutaraldehyde in PBS) and 20 ml of the same ice-cold fixative. Brains were dissected out and post-fixed in the same fixative for 24h at 4°C. The tissues were then washed 3 times with PBS and cryoprotected in 10% [over night (O/N) at 4°C), 20% (O/N at 4°C) and 30% (O/N at 4°C) sucrose in PBS. Tissues were embedded in OCT compound, sectioned at 20 μm, and mounted onto Superfrost plus slides (Thermo Fischer Scientific, Waltham, MA). For in situ hybridization (RNAscope), the manufacturer’s protocol was followed (Advanced Cell Diagnostics, San Francisco, CA). All experiments were replicated in three animals. The probes were designed by the manufacture and available from Advanced Cell Diagnostics. The following probes were used in this study: Mm-Slc17a7-C2 (#416631-C2), Mm-Cck-C1 (#402271-C1), Mm-Tomato-C4 (#317041-C4). Methods of counting and quantifying the cells, and criteria of high level Cck epression. For each animal, Cck expression level was normalized to the average Cck expression in projecting neurons in visual cortex. Neurons expressing low Cck (below the average of Cck expression among projecting neurons in visual cortex) are indicated in grey.

### Associative learning test after CCKBR antagonist application in the AC

After the same anesthesia and surgery as mentioned earlier, a drug infusion cannula was implanted in each hemisphere of the AC of the C57 mouse. The mouse was allowed for recovery for 2 days. For the experimental group, the VS and AS were first presented in pairs to the mouse for 25 trials in each session after the auditory cortex was bilaterally infused with CCKB receptor (CCKBR) antagonist (L365, 260, 10μM in 2% DMSO-ACSF, 0.5μl in both sides, injection speed 0.05μL/min), and for 4 sessions each day during 3 days. For the control group, CCKBR antagonist was replaced with ACSF. On day 4, a baseline test for the percentage of freezing over a time period of 10s was carried out after the VS and AS were presented separately. The AS was then conditioned with footshock for 3 trials. On day 5, post-conditioning tests were carried out to the VS and AS separately. The freezing percentages of different groups to VS were compared by two-way ANOVA.

### Associative learning test after CCK-4 administration in CCK^-/-^ Mice

A drug infusion cannula was implanted on the top of the venous sinus (transverse sinus) of the CCK^-/-^ mouse to administer drugs though i.v. injection. After the i.v. injection of CCK4 (0.01ml, 3.4μM; 1ug/kg) or saline (0.01ml), the VS and AS were immediately presented in pairs to the mouse for 25 trials in each session for 4 sessions each day for 3 consecutive days. The rest of the procedure was the same as the previous experiment. To induce a similar percentage of freezing of C57 mice, the CCK^-/-^ mice needed 9 trials. In a separate experiment to test the dosage of CCK4 that induced panic attack or anxiety, different dosage of CCK4 (2.5, 25 and 250 ug/kg) or vehicle (saline with 5% DMSO) was injected intraperitoneally and the mice activities were monitored for up to 60 minutes after injection. Activities of mice were normalized to the average activities of the vehicle group.

### Histology

After completing all experiments, animal was anesthetized with an overdose of pentobarbital and transcardially perfused with PBS and 4% paraformaldehyde sequentially. Brains were then collected and post-fixed in 4% paraformaldehyde for 24 h. A vibrating blade microtome (VT 1000s, Leica, Germany) was used to cut the brain tissue into sections (60 μm thick). Nissl (Neurotrace 640, 1:200 in 0.01 M PBS with 0.1% Triton X-100, 2h, Thermo Fisher Scientific #N21483, Waltham, MA, USA) or DAPI (1:10000 in 0.01 M PBS, 10 min, Santa Cruz Biotechnology #sc-3598, Dallas, TX, USA) staining was performed in some experiments. Images were obtained with a Nikon Eclipse Ni-E upright fluorescence microscope (Tokyo, Japan) or a Zeiss LSM880 confocal microscope (Oberkochen, Germany).

### Acoustic stimuli, visual stimuli, electrical stimulation, and laser stimulation

All stimuli were generated from computer-controlled RZ6 and RZ5D integrated stimulation stations (Tucker-Davis Technologies [TDT], FL, USA) controlled by self-coded OpenEX program (TDT, FL, USA). Acoustic stimuli were generated as analogue signals and delivered through a close-field speaker (MF1, TDT). The sound pressure level of acoustic stimuli was controlled with OpenEx program and calibrated with a condenser microphone (Center Technology, Taipei). Visual stimuli were generated as analogue signals and delivered though the white LED array. Lasers with wavelengths of 473 nm and 635 nm were generated from different laser generators (Intelligent optogenetic system, Newdoon, China), and 561 nm laser was generated from a separate laser generator (New Industries Optoelectronics Tech Co., China). All lasers were controlled by analogue signals from TDT system. The single pulse width of the laser was always 5 ms.

### Data acquisition and analysis

All *in vivo* electrophysiological data were recorded through TDT system which was controlled by self-coded program in OpenEX (TDT, FL, USA). The filters for spikes and local field potential recordings were set as 500-3,000 Hz and 1-500 Hz, respectively. To identify spikes, the threshold was set as three times the standard deviation of baseline. Offline Sorter (Plexon Inc, US) was used to perform spike sorting, and NeuroExplorer Version 5 (Nex Technologies, US) was used to perform further single unit analysis. Custom Matlab (Mathworks Inc., US) program was used to analyze the recorded fEPSPs. We only included the recording sites in the in vivo or cells in the in vitro experiments with a stable baseline recording. To compare the changes in fEPSP slopes or EPSC amplitudes, the mean of baseline values was first calculated, to which all the slopes or amplitudes were normalized, respectively. For in vivo recording, normalized fEPSP slopes of the last 10 data points (each data point was averaged from the slopes of 6 fEPSPs evoked by VALS, AS or EALS) from baseline and post-pairing test sessions were chosen and averaged to obtain pairs of before and after values for each recording site to perform statistical test. For in vitro patch recording, each data point was averaged from the amplitudes of 3 consecutive EPSCs evoked by VALS or ESAC. And for each cell, the average of 10 baseline data points and the average of last 8 data points of the post-paring session were chosen to generate the pairs of before and after values to perform statistical test. All statistical analyses (paired t-tests, unpaired t-test or two-way RM ANOVA) were performed using SPSS software (IBM, US). Pairwise comparisons were adjusted by Bonferroni correction. Statistical significance was set at p < 0.05.

## Acknowledgements

This work was supported by: Hong Kong Research Grants Council, General Research Fund: 11103220M, 11101521M (GRF, JFH); Hong Kong Research Grants Council, Collaborative Research Fund: C1043-21GF(CRF, JFH); Innovation and Technology Fund: MRP/053/18X, GHP_075_19GD (ITF, JFH); Health and Medical Research Fund: 06172456, 09203656 (HMRF, XC, JFH); The Swedish Research Council: 2018-0273 (TH); The Arvid Carlsson Foundation (TH); The Swedish Research Council: 220-01688 (TH); Key-Area Research and Development Program of Guangdong Province: 2018B030340001(WJS); and the following charitable foundations for their generous support to JFH: Wong Chun Hong Endowed Chair Professorship, Charlie Lee Charitable Foundation, and Fong Shu Fook Tong Foundation.

## Author Contributions

WJS, XC, JFS, TH, and JFH designed the experiments; WJS, YJP, XC and YL collected and analyzed the data of the physiological part; HHW and MZ collected and analyzed the RNAscope data; YJP, XJZ and HMF collected and analyzed the data of the behavioral part; WJS, PT, HL and JL performed viral injection, retrograde tracing, and IHC experiments; WJS, XC, TH and JFH wrote the manuscript.

---

**Supplementary Figure 1.**
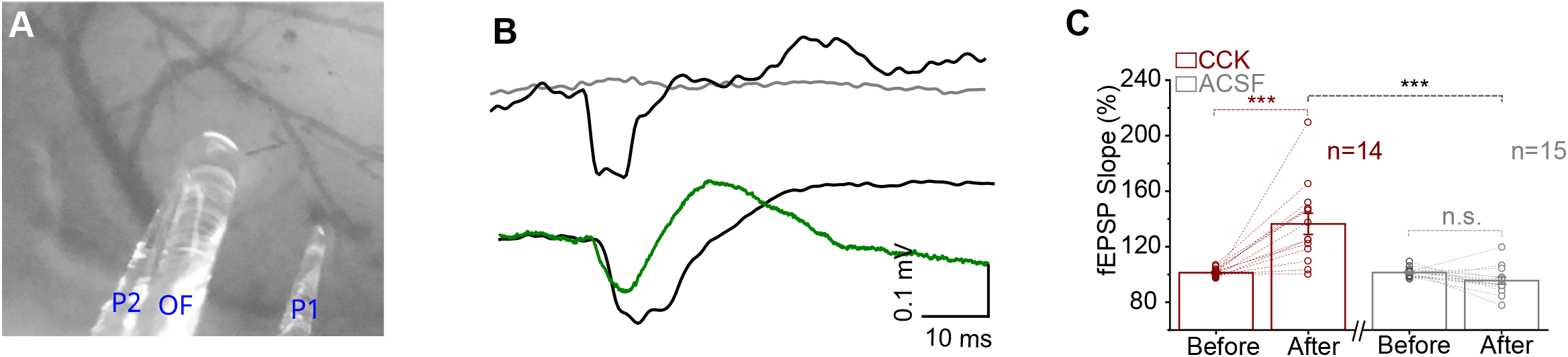
(A) An example image of the positions of glass pipette recording electrodes (P1 and P2) and optical fiber (OF) for laser stimulation. (B) Upper: example fEPSP traces recorded by glass (Gray) or tungsten (black, artefact) electrode in wild type mice without any expression in the recording site. Bottom, example fEPSP traces recorded by glass (green, real signal) or tungsten (black, mixture of artefact and true signal) electrodes in the Cck-iRES-Cre mice with the expression of ChR2 in the projecting terminals of the recording area. (C) Individual and average fEPSPVALS slope changes before and after Pre/Post Pairing with CCK-8S (red) or ACSF (gray) infusion in the AC. *** p < 0.001, n.s. p = 0.411, n = 14 for CCK group, n = 15 for ACSF group, two-way RM ANOVA with post-hoc Bonferroni test. See Table S1 for detailed Statistics.

**Supplementary Figure 2.**
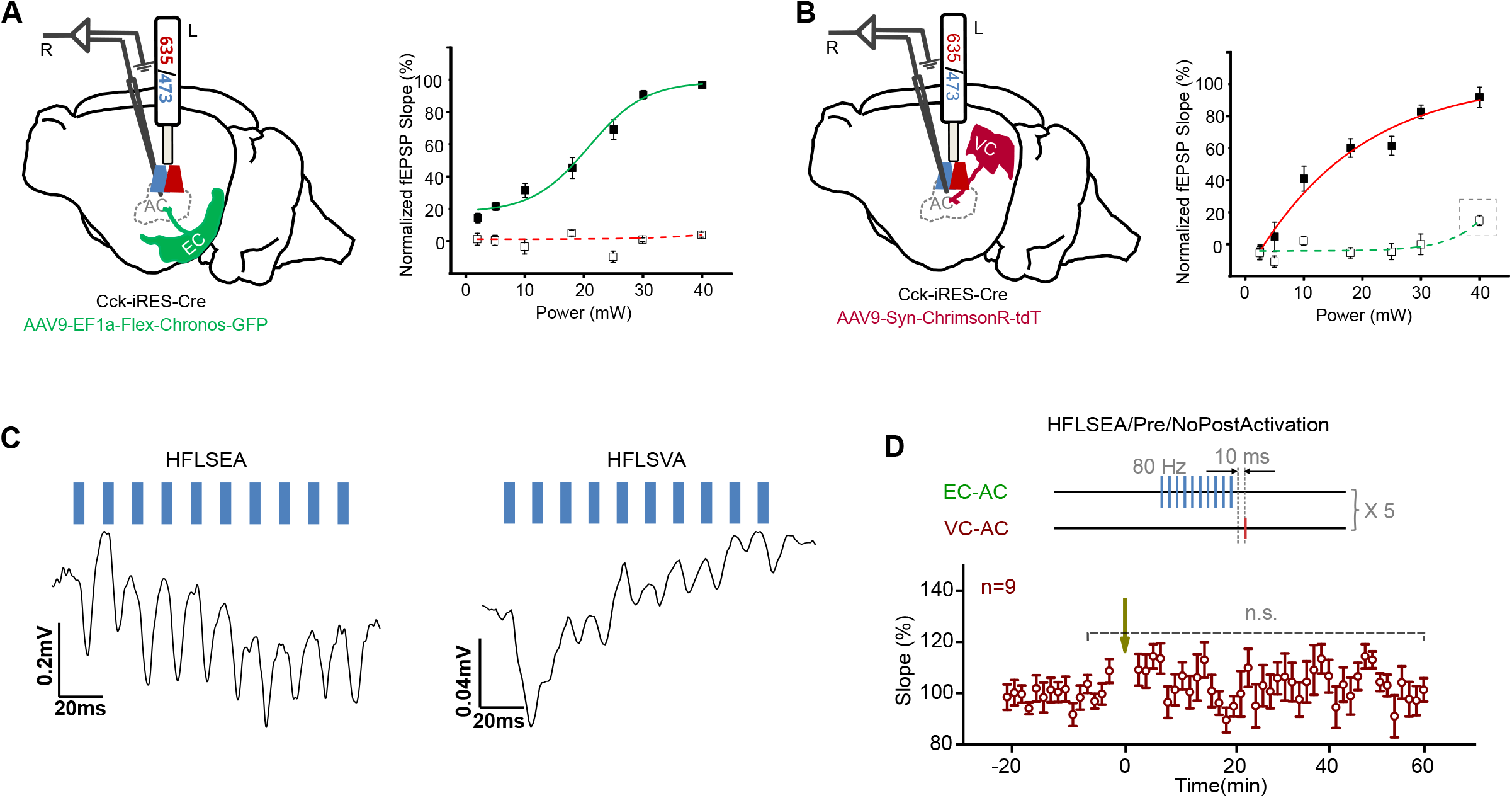
(A) and (B) Optimal laser power determination of 473 nm and 635 nm to prevent crosstalk. Diagram of the experimental design, animals with only one injection of either AAV9-Ef1α-Flex-Chronos-GFP into the entorhinal cortex (A left) or AAV9-Syn-ChrimsonR-tdTomato into the visual cortex (B left) were prepared, 473 nm and 635 nm were both applied to stimulate their terminals in the auditory cortex, and their corresponding local field potentials were recorded by glass pipette electrodes. A right, normalized fEPSPs slopes (n=9) evoked by 473 nm (green line) or 635 nm (red dashed line) laser stimulation of CCK+ ENT→AC projection terminals; B right, normalized fEPSPs slopes (n=12) evoked by 473 nm (green dashed line) or 635 nm (red solid line) laser stimulation of VC→AC universal projection terminals; gray dashed rectangle in B-right emphasizes the power that may induce crosstalk. (B) Example traces of HFLSEA (left) and HFLSVA (right). Scale bars: left, 20 ms and 0.2 mV; right, 20 ms and 0.04 mV. (C) Upper: schematic drawing of the protocol of HFLSEA/Pre/NoPostActivation. Bottom: normalized fEPSPVALS slopes before and after HFLSEA/Pre/NoPostActivation. Error bars represent SEM. paired t-test, t (8) = -0.899, n.s. p = 0.395, n = 9. See Table S1 for detailed Statistics.

**Supplementary Figure 3.**
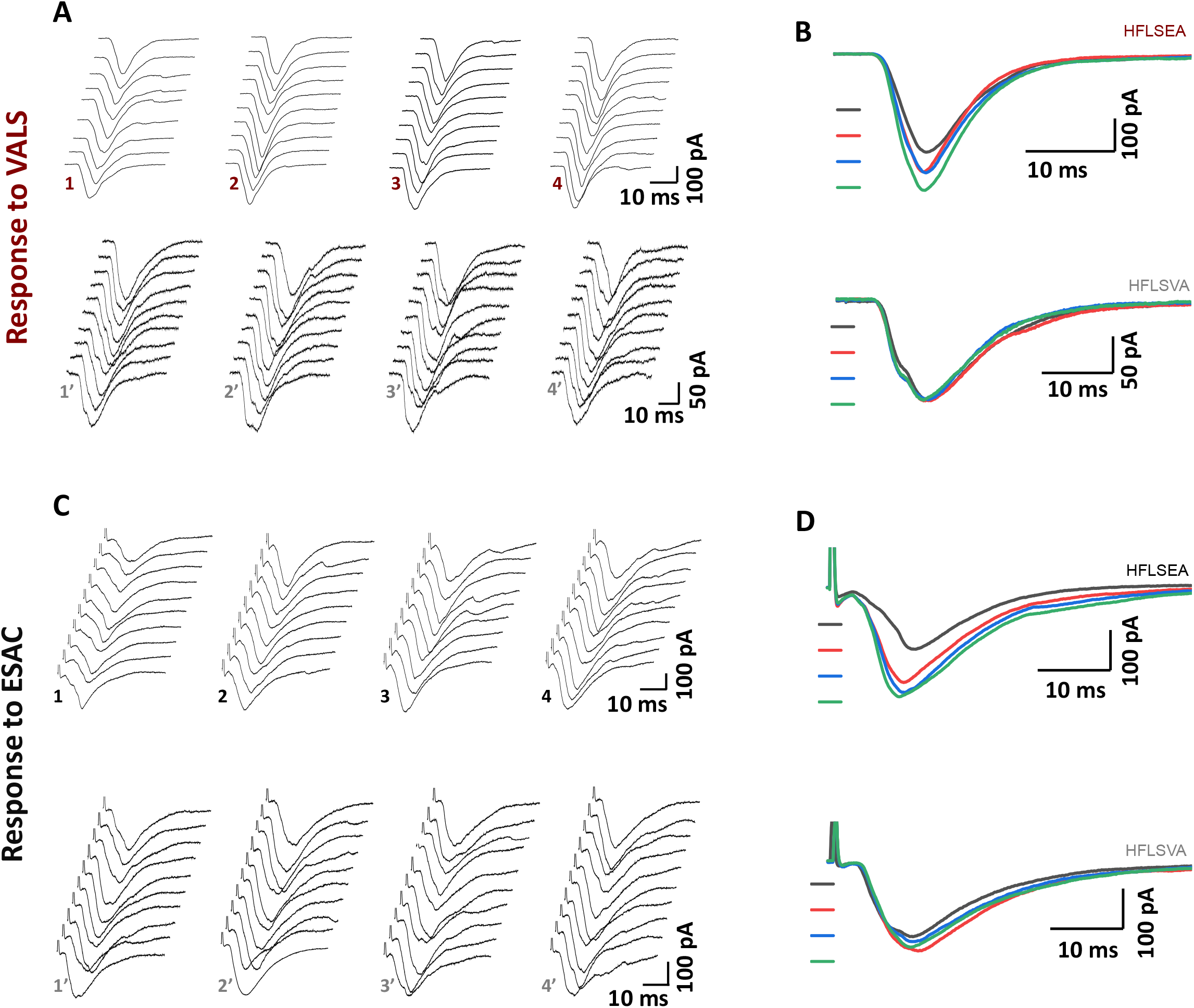
(A) Representative EPSCVALS traces at different timepoints and (B) their averaged traces in the HFLSEA/Pre/PostPairing (upper) or HFLSVA/Pre/PostPairing (bottom) group. (1 or 1’: first 10 consecutive individual trace before pairing; 2-4 or 2’-4’: 10 consecutive individual trace 0 min, 13 min and 27 min after pairing respectively). (C) Representative EPSCESAC traces at different timepoints and (D) their averaged traces in the HFLSEA/Pre/PostPairing (upper) or HFLSVA/Pre/PostPairing (bottom) group. (1 or 1’: first 10 consecutive individual trace before pairing; 2-4 or 2’-4’: 10 consecutive individual trace 0 min, 13 min and 27 min after pairing respectively). See Table S1 for detailed Statistics.

**Supplementary Figure 4.**
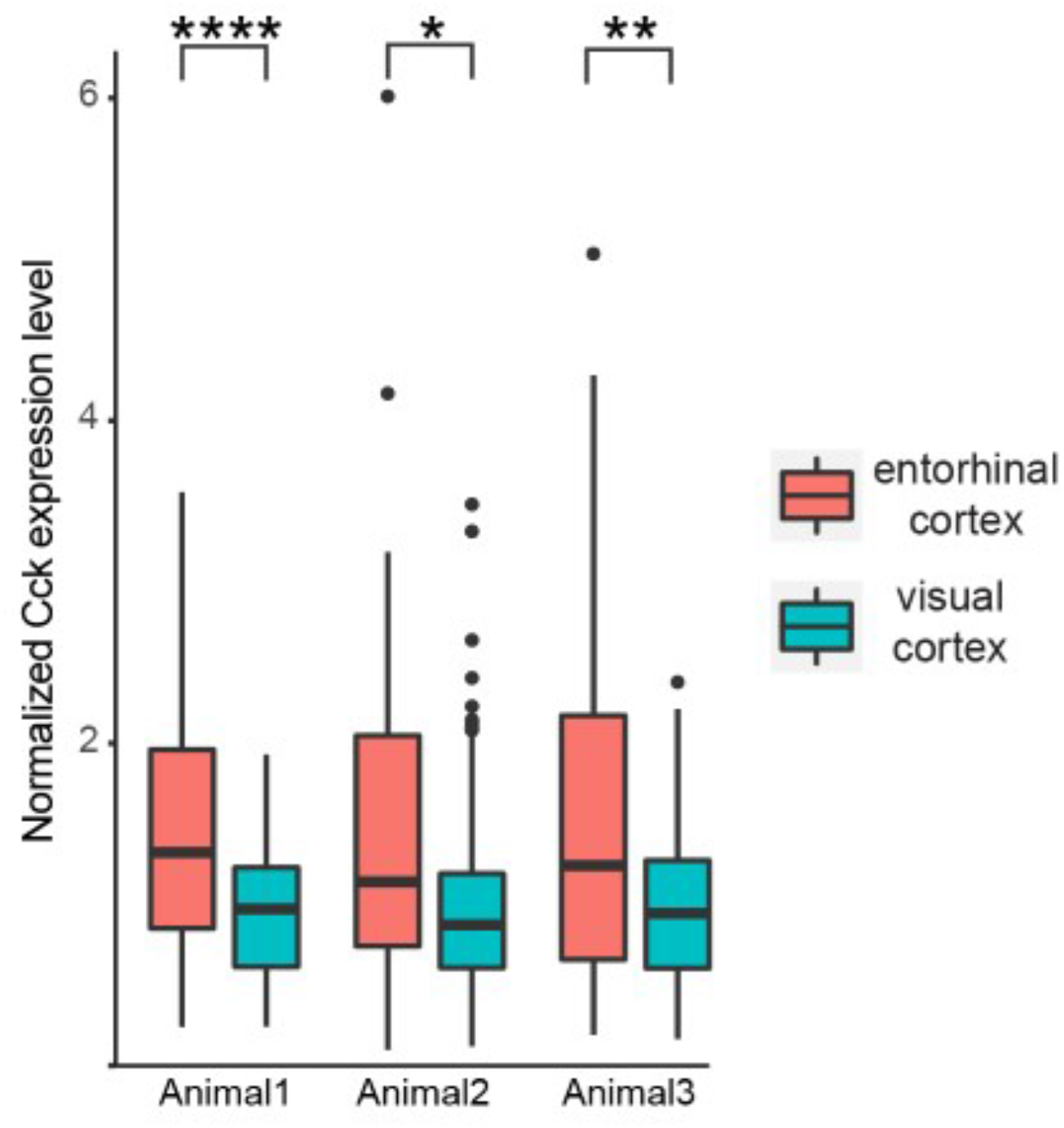
Comparison of Cck expression level in neurons in the Ent and VC which project to the VC. For each animal, Cck expression level is normalized to the average Cck expression in projecting neurons in visual cortex. Unpaired t-test, ****p<0.0001, **p<0.01, *p<0.05. See Table S1 for detailed Statistics.

**Supplementary Figure 5.**
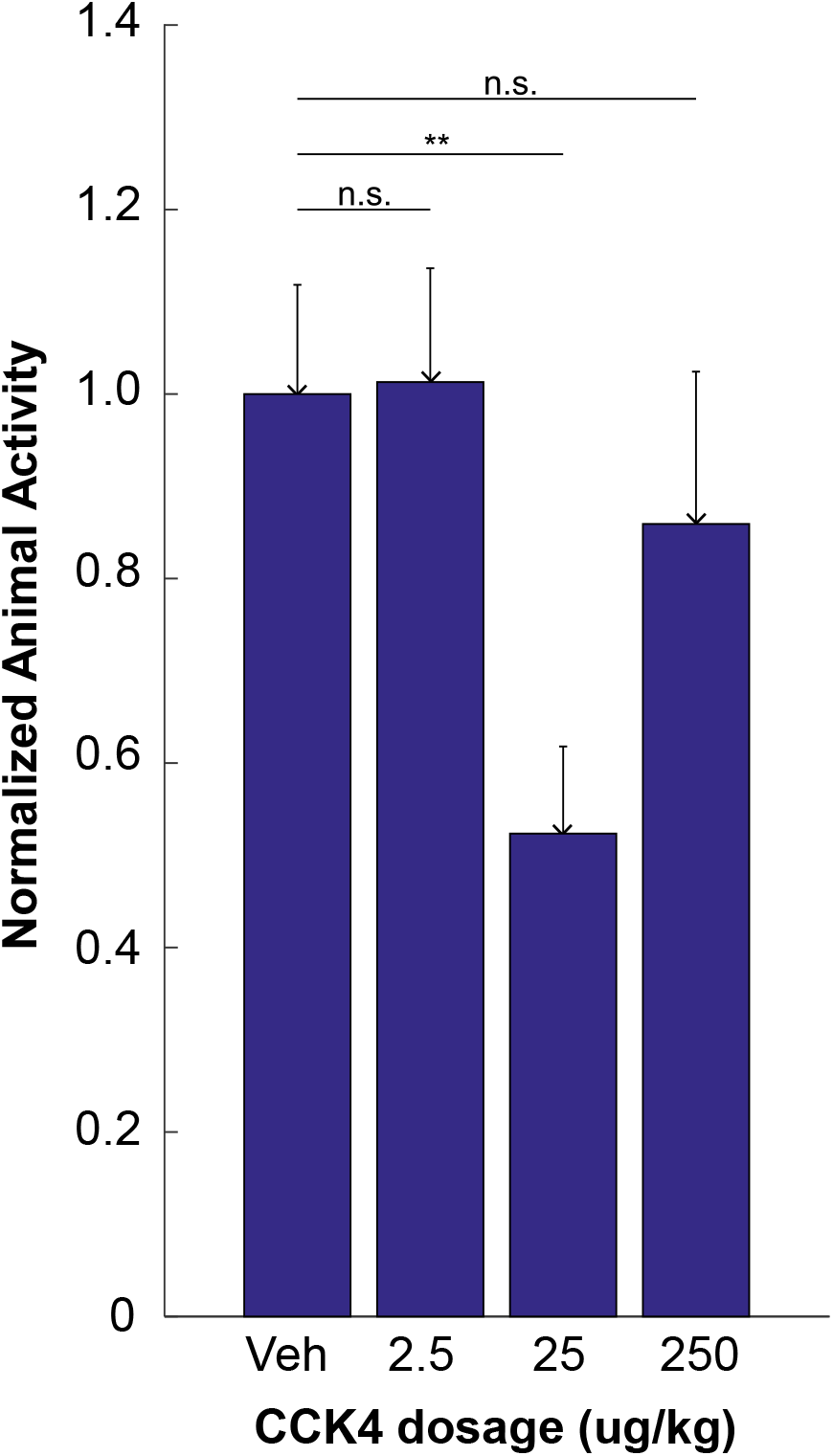
Animal activities after different dosage of CCK4 injection. Data shows normalized activities of mice since 60 seconds after CCK4 or vehicle injection up until 180s. n = 10 for each group. **, p<0.01; n.s., not significant; one-way ANOVA.

**Table S1.**
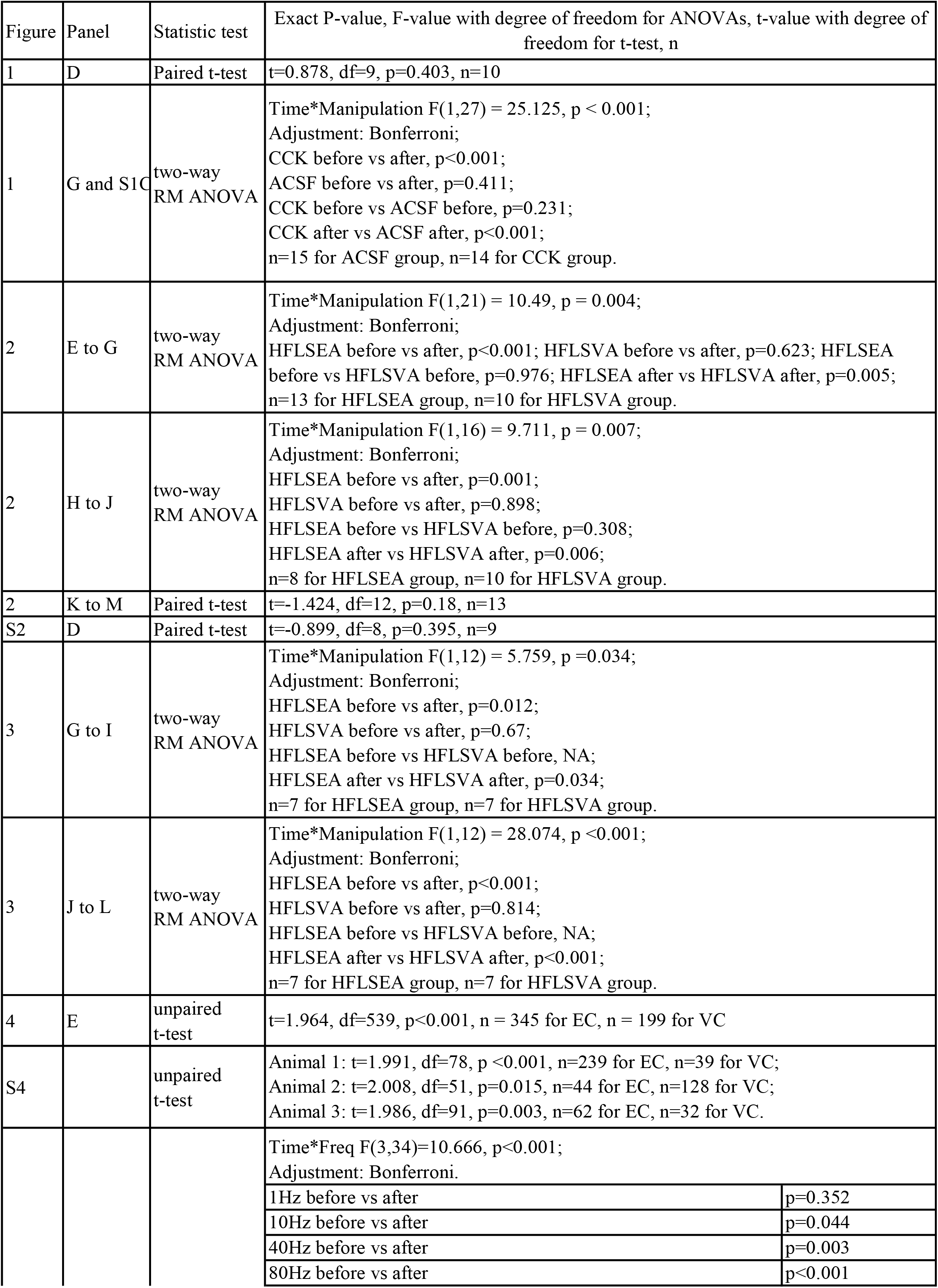
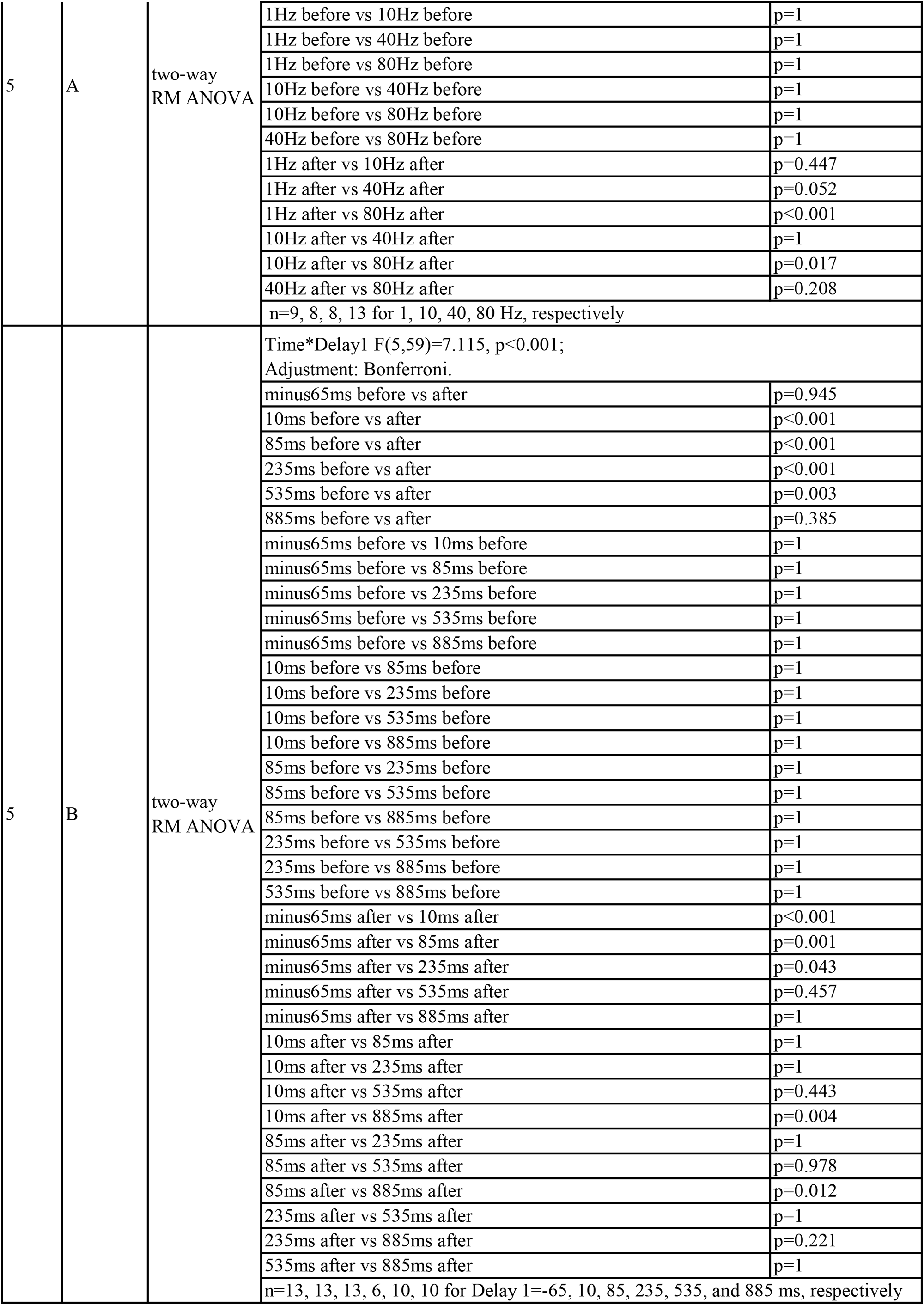
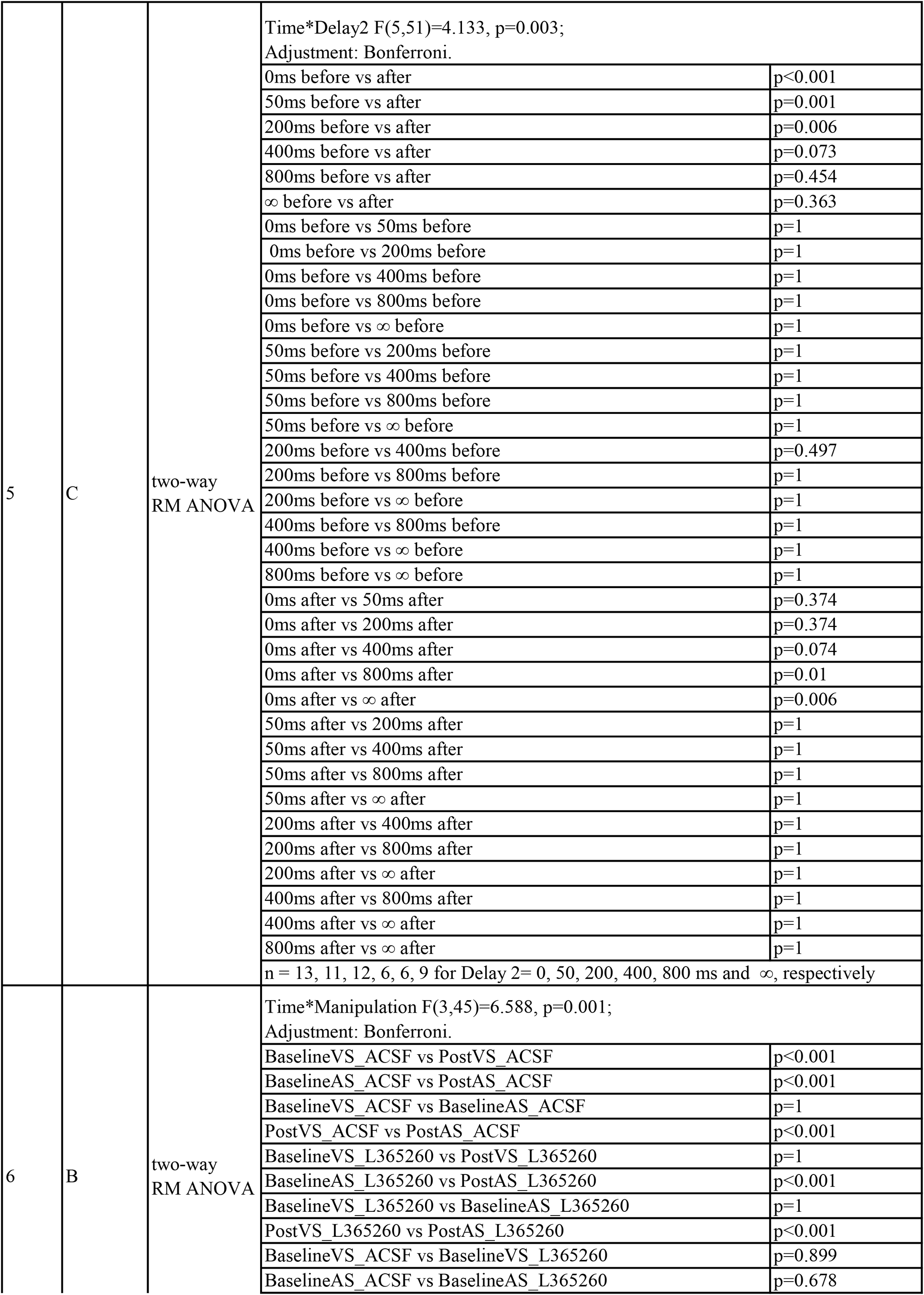
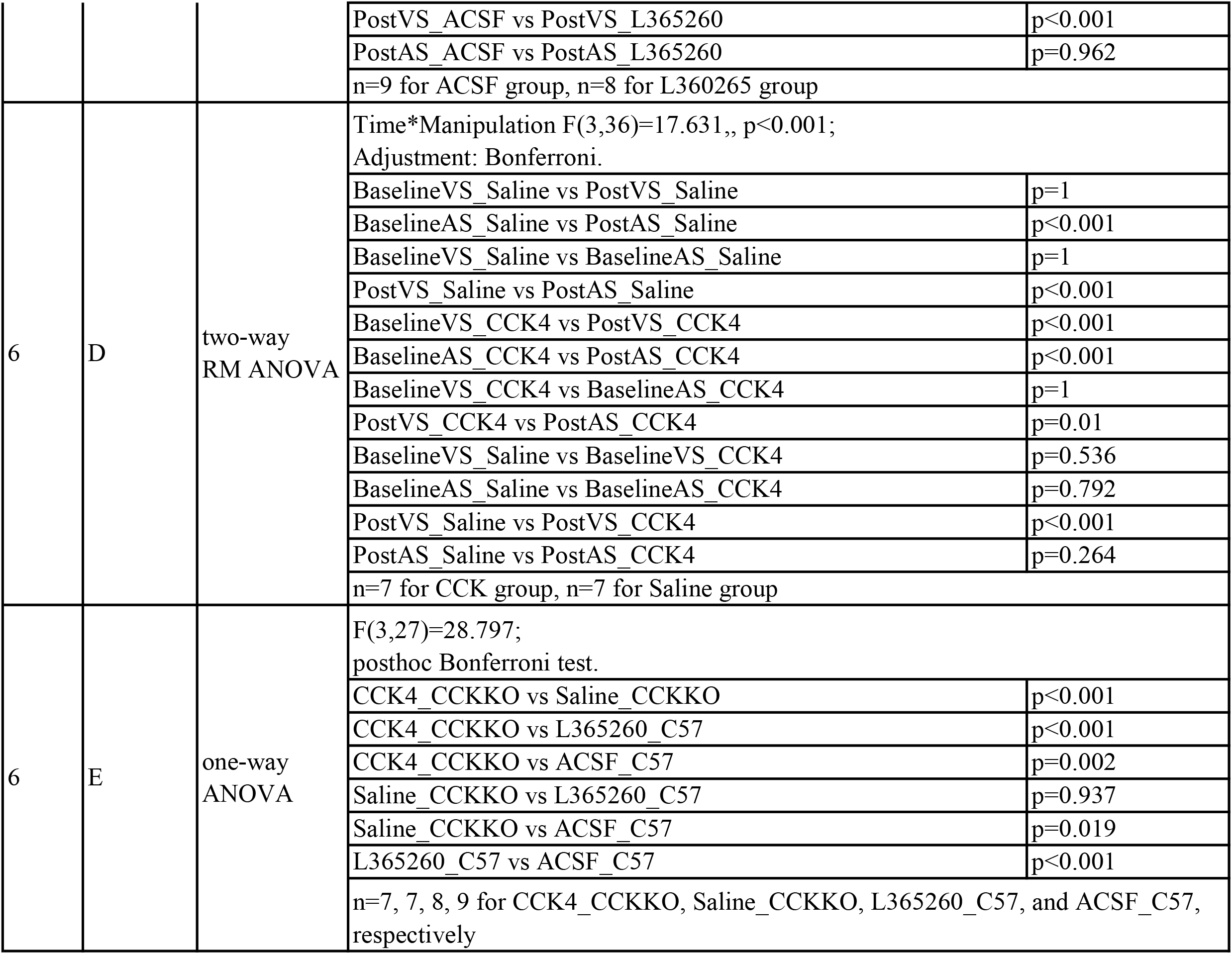
Statistics

